# A TauP301L mouse model of dementia; development of pathology, synaptic transmission, microglial response and cognition throughout life

**DOI:** 10.1101/420398

**Authors:** Zelah Joel, Pablo Izquierdo, Wenfei Liu, Chloe Hall, Martha Roberts, Kenrick Yap, Amy Nick, Emma Spowart, Roshni Desai, Laurenz Muessig, Rivka Steinberg, Stuart Martin, Kenneth J Smith, Jill C Richardson, Francesca Cacucci, Dervis A. Salih, Damian M. Cummings, Frances A. Edwards

**Affiliations:** University College London, Gower Street, London, WC1E 6BT, UK

## Abstract

**Background:** Late stage Alzheimer’s disease and other dementias are associated with neurofibrillary tangles and neurodegeneration. Here we describe a mouse (TauD35) carrying human Tau with the P301L mutation that results in Tau hyperphosphorylation and tangles. Previously we have compared gene expression in TauD35 mice to mice which develop plaques but no tangles. A similar comparison of other pathological features throughout disease progression is made here between amyloidβ and Tau mice described in Parts I and II of this study.

**Methods:** *In vitro* CA1 patch clamp and field recordings were used to investigate synaptic transmission and plasticity. Plaque load and microglia were investigated with immunohistochemistry. Cognition, locomotor activity and anxiety-related behaviours were assessed with a forced-alternation T-maze, open field and light/dark box.

**Results:** Transgene copy number in TauD35 mice fell into two groups (HighTAU and LowTAU), allowing assessment of dose-dependent effects of overexpression and resulting in tangle load increasing 100-fold for a 2-fold change in protein levels. Tangles were first detected at 8 (HighTAU) or 13 months (LowTAU) but the effects on synaptic transmission and plasticity and behaviour were subtle. However, severe neurodegeneration occurred in HighTAU mice at around 17 months, preceded by considerable proliferation and activation of microglia at 13 months of age; both increasing further at 17 months. LowTAU mice at 24 months of age showed a comparable tangle load and microglial proliferation to that occurring at 13 months in HighTAU mice. However, LowTAU mice showed no neurodegeneration at this stage and considerable microglial activation, stressing the dependence of these effects on overexpression and/or age.

**Conclusions:** Comparison of the effects of amyloidβ and plaques without tangles in a model of preclinical Alzheimer’s disease to the effects of tangles without amyloidβ plaques in the late stage model described here may clarify the progressive stages of Alzheimer’s disease. While Tau hyperphosphorylation and neurofibrillary tangles are eventually sufficient to cause severe neurodegeneration, initial effects on synaptic transmission and the immune response are subtle. In contrast while even with a heavy plaque load little if any neurodegeneration occurs, considerable effects on synaptic transmission and the immune system result, even before plaques are detectable.

## Background

The prevalence of dementia in the ageing population is becoming a major problem for the health care system. Understanding the progression of the disease and the links between pathology and cognitive outcomes is essential if we are to find ways of slowing or preventing the onset of dementia. Alzheimer’s disease is the most common form of dementia and, in its later stages, shares common pathologies with frontotemporal dementia and other tauopathies. In all these dementias, the development of neurodegeneration is closely correlated with the hyperphosphorylation of the microtubule associated protein Tau and development of neurofibrillary tangles. In the case of Alzheimer’s disease this development is subsequent to a rise in amyloidβ (Aβ) and deposition of plaques, whereas some of the other tauopathies occur as a direct result of mutations in the Tau gene.

To study the relative time course of synaptic and microglial changes and neurodegeneration in relation to phosphorylation of Tau and neurofibrillary tangles, we have characterised a novel strain of mice (TauD35), transgenic for *MAPT*, which encodes human Tau protein with a mutation that, in man, results in frontotemporal dementia with parkinsonism linked to chromosome 17 ^[1]^. The pathology in this mouse progresses much more slowly than previously described models of Tauopathy ^[2, 3]^. Previously we have correlated gene expression and development of neurofibrillary tangles in this mouse and undertaken gene expression network analysis finding that most gene expression changes, including upregulation of microglial genes and decrease in expression of synaptic genes, occur only very late in disease progression, months after neurofibrillary tangles become evident ^[1]^. This is in contrast to more aggressive overexpression models in which neurodegeneration occurs at a young age and synaptic changes are seen even before neurofibrillary tangles are evident ^[4]^. A very interesting study in another Tau model showed synaptic and behavioural changes in middle age (approximately 10 months) which were reversible on turning off the transgene ^[5, 6]^. However, further work is required to elucidate the mechanisms of this progression ^[7]^. It is thought that neurofibrillary tangles may underlie neuronal death by interfering with axonal transport but mitochondrial dysfunction, leading to calcium homeostasis breakdown and apoptosis, likely also contributes ^[7]^. Evidence suggests that caspases are involved, especially at earlier stages ^[8-10]^ in these aggressive models but in the slower progressing model TauD35 mice, while some caspases show increased expression, this again occurs only late in disease progression ^[1; www.mouseac.org]^.

The more aggressive models, such as rTg4510, are useful in that they start to develop neurofibrillary tangles very early and show rapid neurodegeneration, which is convenient in terms of cost and various practical considerations and interesting data have emerged ^[2, 3]^. However, it makes the separation of stages of disease progression difficult and means that it occurs in mice at ages equivalent to adolescence through to early middle age rather than old age, as is most common in Alzheimer’s disease. In the present study we characterised mice developed at GlaxoSmithKline (TauD35) with the MAPT_P301L_ mutation also on the CamKIIα promoter (the same mutation and promoter as for the rTg4510 mice but with much slower progression of pathology). During this study we established that the TauD35 line in fact consisted of two lines of these mice with different copy numbers (referred to here as LowTAU and HighTAU) enabling us to assess the dependency of changes seen on the gene dosage, albeit *post hoc*.

## Methods

Methods are the same as for ^[11]^ except adapted for neurofibrillary tangles.

### Animals

All experiments were performed in agreement with the Animals (Scientific Procedures) Act 1986, with local ethical approval and in agreement with the GlaxoSmithKline statement on use of animals.

Transgenic TauD35 male mice were generated by GlaxoSmithKline (Harlow, UK) on the background mouse line C57Bl/6J (Charles River, UK) via pronuclear injection and bred and maintained at Charles River Laboratories. The mice harbour human cDNA for the 0N4R isoform of MAPT carrying the P301L mutation under the alpha isoform of the Ca2+/calmodulin dependent protein kinase II (CaMKII) promoter. Age-matched wild type littermates were used as controls.

Mice from Charles River were shipped to UCL at 3-months-old. Mice were kept in large cages (20 x 35 x 45 cm) with an enriched environment. Thus, cages containing 2-8 male mice (TauD35 and littermate wild type controls) were maintained in a 12-hour light/12-hour dark cycle with food and water *ad libitum*. Environmental enrichment consisted of changes of food location, bedding type (e.g. tissue, shredded paper, paper roll, paper bags) and inanimate objects (e.g. running wheels, rodent balls, tubing, houses (mostly purchased from Eli Lilly Holdings Limited, Basingstoke, UK)) within the cage at least once per week. Mice were used for experimentation at the ages stated (± 0.5 months) and, where unavoidable, were single-housed for no longer than 24 hours. Tails or ear punches were genotyped using standard PCR protocols.

### Genotyping

#### Genotype confirmation using conventional PCR methods

Briefly, genomic DNA was extracted using the ‘HotSHOT’ lysis method. Alkaline lysis reagent (25 mM NaOH, 0.2 mM EDTA, pH12) was added to tissue samples prior to heating to 95°C for 30 minutes. The sample was then cooled to 4°C before the addition of neutralisation buffer (40 mM Tris-HCl, pH 5). The PCR reaction was performed through addition of MyTaq DNA Polymerase (Bioline) reaction buffer and primer pair:

5’-AAGACCAAGAGGGTGACACGG-3’,

5’-CCCGTCTTTGCTTTTACTGACC-3’

using the cycling parameters: 94°C (2 minutes), 58°C (30 s), 72°C (30 s), for 30 cycles and a final extension at 72°C for 4 minutes. PCR product size 130 bp.

#### Transgene copy number confirmation

Genomic DNA was extracted using the ‘HotSHOT’ lysis method described above. Unique TaqMan primer and probe sequences were generated using Applied Biosystems custom design sequence using the published sequence for the CaMKIIα promoter sequence (Accession# AJ222796). The qPCR assay was run with 4 μl of genomic DNA per well using the CaMKIIP_CCVI3TR primer and probe set (catalogue number 4400294, Applied Biosystems), using the Taqman genotyping master mix with each sample run in triplicate in parallel with the Taqman copy number reference assay for mouse Tfrc (catalogue number 4458370, Applied Biosystems), using the manufacturer’s instructions.

### Western blot

#### Brain tissue extraction

Brains were removed from the skull on ice and hippocampus extracted within 5 minutes of death, then snap frozen on dry ice and stored at -80°C until protein extraction was performed.

#### Protein extraction

Mouse hippocampal samples were sonicated (Branson Sonifier, 450) for 30 s at 9-12 W in RIPA buffer (1% [v/v] Triton X-100, 1% [w/v] sodium deoxycholate, 0.1% [w/v] SDS, 0.15 M NaCl, 20 mM Tris-HCl (pH 7.4), 2 mM EDTA (pH 8.0), 50 mM NaF, 40 mM β- glycerophosphate, 1 mM EGTA (pH8.0), 2 mM sodium orthovanadate, protease inhibitor tablet (cat no 11 836 153 001; Roche), aprotinin and phosphatase inhibitor cocktail 1 and 2 (1:100; Sigma), 1 mM PMSF: approximately 1 ml for 100 mg tissue).

For total protein extracts, homogenates were spun at 18,000 g for 10 minutes at 4°C and the supernatant collected. Sample buffer (312.5 mM Tris (pH 6.8), 10% [w/v] SDS, 250 mM DTT, 50% [v/v] glycerol, 0.025% [w/v] bromophenol blue) was added prior to boiling the samples at 95°C for 10 minutes and storing at -20°C.

Sarkosyl-insoluble tau extracts were obtained as described previously ^[3]^ 90µl of homogenate was ultracentrifuged at 150,000g for 15 minutes at 4°C. The supernatant (S1) was removed for storage and the pellet re-homogenised in 4 volumes of 10mM Tris (pH 7.4), 0.8M NaCl, 1mM EGTA, 10% sucrose, 1mM PMSF and ultracentrifuged as above. The resulting pellet was discarded and the supernatant incubated with 1% sarkosyl (30% sarkosyl NL30, BDH) for 1hr at 37°C, prior to ultracentrifugation at 150,000g for 30 minutes at 4°C. The supernatant was again collected for storage (S2) and the pellet re-suspended in 20µl 10mM Tris (pH 8) and 1mM EDTA and labelled P3: sarkosyl insoluble tau. Sample buffer (312.5mM Tris (pH 6.8), 10% [w/v] SDS, 250mM DTT, 50% [v/v] glycerol, 0.025% [w/v] bromophenol blue) was added to all fractions prior to boiling the samples at 95°C for 10 minutes and storing at -20°C.

#### Western blot analysis

Protein corrected (Bradford assay) samples were resolved by sodium dodecyl sulphate-polyacrylamide gel electrophoresis (SDS-PAGE) using 8-10% polyacrylamide gels. Molecular weights were verified using a molecular weight standard (BioRad, #161-0374). Proteins were transferred to a nitrocellulose membrane (0.45 μm, BioRad) using electroblotting (30 V, overnight). Following transfer, membranes were washed in Tris-buffered saline (Tris), prior to blocking against non-specific antibody binding in 5% milk/Tris for 1 hour at room temperature. Membranes were incubated in primary antibody in 5% milk/Tris overnight at 4°C. Following overnight incubation, membranes were washed in 0.05% Tween-20/Tris for five changes of 7 minutes each at room temperature and then for 10 minutes in 5% milk/Tris. Horseradish peroxidase-conjugated goat anti-mouse (1:10,000; Jackson ImmunoResearch, #115-035-146) incubation was performed in 5% milk/Tris for 1 hour at room temperature. Membranes were washed again for five changes of 7 minutes in 0.05% Tween-20/Tris at room temperature, before a final wash of 5 minutes in Tris. Peroxidase activity was revealed using an enhanced chemiluminescence detection kit (ECL, Amersham). Image acquisition and densitometric analysis was performed using ImageLab (v4.1, BioRad) as described ^[12]^. Density values for bands of interest were normalized against heat shock protein 9 (HspA9, GRP75 (N52A/42) Cambridge Biosciences, #MMS-5164-100).

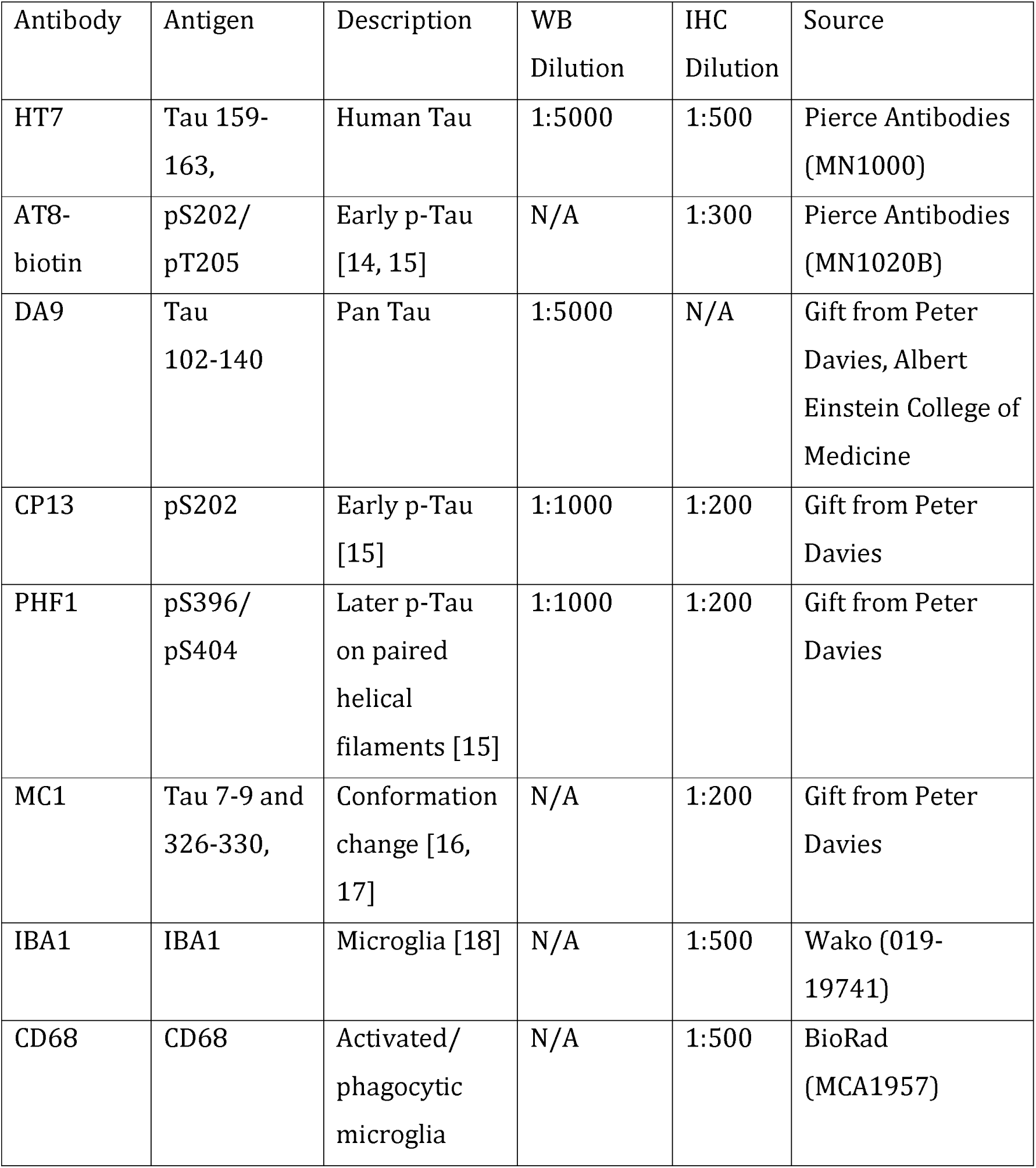
Antibodies used for western blots and immunohistochemistry: (for review see ^[13]^)

### Immunohistochemistry

Animals were deeply anaesthetised (1:10 Euthatal:Intra-Epicaine, National Veterinary Supplies) and transcardially perfused with 0.1 M phosphate buffer saline (PBS) followed by 10% buffered formal saline (Pioneer Research Chemicals Ltd). Alternatively, single hemispheres were drop-fixed immediately following brain extraction for electrophysiology. The brains were post-fixed in 10% buffered formal saline for 24hrs and cryoprotected in 30% sucrose/0.03% sodium azide/PBS at 4 C for at least 24hrs before sectioning or storage. Transverse sections were cut at 30 μm through the full left hippocampus using a frozen sledge microtome (SM 2000 R, Leica) and collected into a 24-well plate containing PBS/sodium azide (0.03%) for storage at 4°C. Serial sections were placed in separate wells until all wells contained a section and collection then continued serially from Well 1 so that within each well the transverse sections were from the length of the hippocampus at least 720µm apart.

Standard immunohistochemistry techniques were employed. Sections for all immunohistochemistry were washed in PBS, followed by 0.3% Triton X-100 in PBS (PBST) and subsequent blocking in 8% horse serum/PBST for 1 hour. Incubation with primary antibody in blocking solution was performed overnight at 4°C. Sections were again washed with PBST. The appropriate Alexa-conjugated secondary antibody (1:500; Invitrogen) was added to blocking solution for a 2-hour incubation at room temperature in the dark. Following PBS wash, DAPI was applied to all sections for 5 minutes. Sections were washed for a final time in PBS before mounting. Sections from age-matched wild type control mice were stained in parallel for all ages. Sections were mounted in anatomical order onto SuperFrost Plus glass slides by floating on PBS and then cover-slipped using Fluoromount G mounting medium.

#### Nissl Stain

Sections were mounted and left to dry overnight prior to staining. Sections were dipped in H_2_0 and then submerged in 1% w/v cresyl violet (Alfa Aesar, Massachusetts) for 2 minutes and blotted to remove excess solution. Sections were de-stained in 70% ethanol containing 1% glacial acetic acid for <1 minute and then dried by submersion in 100% ethanol for 2 minutes followed by submersion in xylene (3 minutes) to completely remove water. Sections were then cover-slipped using DPX mounting medium.

#### Imaging and data analysis

Sections were imaged for quantification using an EVOS FL Auto Cell Imaging System (Life technologies). For immunohistochemistry and Nissl staining, tiled images were taken of the whole (transverse) hippocampus using a 20X objective. For immunohistochemistry and cell counts an area of 400 μm x 240 μm was defined in the CA1 and CA3. Cell counts were performed using Adobe Photoshop CS6. Iba1-positive microglia were only counted if a DAPI-positive nucleus was also present. Neurofibrillary tangle counts for double labelled sections (AT8/HT7) included only cells positive for both antibodies. Cell counts using Nissl staining included all principle cells within the defined area, with cells identified by their morphology. Due to high density Nissl staining in the dentate gyrus, cell counts could not be reliably obtained for this area. Where neurofibrillary tangles/cells could not be individually distinguished, but stain was present, a count of 1 was awarded. Neurofibrillary tangle and cell counts are therefore an underestimate of true number. Sample sizes reported for immunohistochemistry and cell counts indicate the number of animals used to calculate means. A minimum of 3 sections were used to create a mean for each animal. Sections for any given condition were obtained from the same collection well within the 24-well plate and were therefore a minimum of 720 μm apart, thus avoiding multiple counts of the same cells. Confocal images were taken using a laser scanning confocal microscope (Zeiss LSM 510) with a 60X Plan-Apochromat oil immersion objective (1.4 numerical aperture).

### Electrophysiological recordings

#### Acute hippocampal brain slice preparation

Mice were decapitated and the brain rapidly removed and placed in ice-cold dissection artificial cerebrospinal fluid (ACSF, containing (in mM): 125 NaCl, 2.4 KCl, 26 NaHCO_3_, 1.4 NaH_2_PO_4_, 20 D-glucose, 3 MgCl_2_, 0.5 CaCl_2_, pH 7.4, ~315 mOsm/l). After approximately two minutes in ice-cold dissection ACSF, the brain was prepared for slicing by removing the cerebellum, hemisection of the forebrain and a segment cut away from the dorsal aspect of each hemisphere at an angle of approximately 110° from the midline surface to optimise slicing transverse to the hippocampus. Each hemisphere was then glued with cyanoacrylate (Loctite 406, Henkel Loctite Limited, UK) on this surface onto the stage of a vibrating microtome (Integraslice model 7550 MM, Campden Instruments, Loughborough, UK) containing frozen dissection ACSF and 400 μm transverse slices of hippocampus cut. As each slice of a hemisphere was cut, the hippocampus was dissected out, retaining a portion of entorhinal cortex and the resulting smaller slice was placed into a chamber containing ‘Carbogenated’ (95% O_2_/5% CO_2_; BOC Limited) dissection ACSF at room temperature (approximately 21°C). After 5 minutes, slices were then transferred into a fresh chamber held at 36°C with the same dissection ACSF. At 5-minute intervals, they were then consecutively transferred to physiological Ca^2+^ and Mg^2+^ ion concentrations (in mM): i) 1 Mg^2+^, 0.5 Ca^2+^; ii) 1 Mg^2+^, 1 Ca^2+^; iii) 1Mg^2+^, 2 Ca^2+^. After approximately 20 minutes at 35°C (i.e., once transferred into the 1 Mg^2+^, 2 Ca^2+^ ACSF).

#### Patch-clamp recordings in brain slices

Once transferred to 1 Mg^2+^, 2 Ca^2+^ ACSF, slices were allowed to return to room temperature and after at least a further 40 minutes recovery time, a single slice was transferred to a submerged chamber and superfused with recording ACSF (containing (in mM): 125 NaCl, 2.4 KCl, 26 NaHCO_3_, 1.4 NaH_2_PO_4_, 20 D-glucose, 1 MgCl_2_, 2 CaCl_2_, bubbled with Carbogen). Individual CA1 pyramidal or dentate gyrus granule neurones were visualised using infrared-differential interference contrast microscopy on an upright microscope (model BX50WI, Olympus, UK). Glass microelectrodes for patch-clamp were pulled from borosilicate glass capillaries (Catalogue number GC150F-7.5, 1.5 mm OD x 0.86 mm ID, Biochrom-Harvard Apparatus Ltd, Cambridge, UK) on a vertical puller (model PP830, Narishige International Ltd, London UK). Electrodes (tip resistance approximately 5 MΩ) were filled with a CsCl-based internal solution (containing (in mM): CsCl 140, HEPES 5, EGTA 10, Mg-ATP 2, pH 7.4, ~290 mOsm/l). Patch-clamp recordings were performed using an Axopatch 1D (Molecular Devices, Sunyvale, CA, USA), and current signals low-pass filtered at 10 kHz then 2 kHz (Brownlee Precision Model 440, NeuroPhase, Santa Clara, CA, USA) during digitization (10 kHz; 1401plus, Cambridge Electronic Design, Limited, Cambridge, UK) and acquired using WinWCP (for isolated events; version 4.6.1; John Dempster, Strathclyde University, UK) and WinEDR (for continuous recordings; John Dempster, Strathclyde University, UK). Stimulation was applied *via* a patch electrode filled with ACSF, placed extracellularly in the appropriate axon path using a square pulse constant-voltage stimulator (100 µs; DS2A-MkII, Digitimer Ltd, UK) triggered by WinWCP.

WinEDR synaptic analysis software was used for detection of spontaneous and miniature currents and WinWCP used to analyse identified spontaneous, miniature and evoked currents. Criteria for detection of spontaneous or miniature currents was to remain over a threshold of 3 pA for 2 ms. Currents were inspected by eye and only included if the rise time was <3 ms and faster than the decay.

#### Field potential recordings in brain slices

Slices were transferred as needed to a heated (30±1°C) submerged chamber and superfused with ACSF and allowed to recover for 1 h in the recording chamber. A glass stimulating electrode (filled with ACSF, resistance ~2 MΩ) and an identical recording electrode (connected to an AxoClamp 1B via a 1X gain headstage) were both positioned in stratum radiatum of the CA1 field to obtain a dendritic excitatory postsynaptic field potential (fEPSP). Recordings were controlled and recorded using WinWCP software (as above), filtered at 10 kHz and subsequently at 3 kHz and digitized at 10 kHz via a micro1401 interface (Cambridge Electrical Designs, UK). Stimuli (constant voltage 10-70V, 100 μs; model Digitimer DS2A-MkII or Grass SD9) were applied at 0.1 Hz and resultant fEPSPs subsequently averaged over consecutive 1-minute intervals. Stimulation intensity was set at approximately 30-50% of the intensity required to evoke a population spike or the maximum fEPSP amplitude obtained and a ≥15-minute stable baseline recorded. LTP conditioning was applied at test-pulse stimulus intensity and consisted of 3 trains of tetani, each consisting of 20 pulses at 100 Hz, 1.5 s inter-train interval. Following conditioning, fEPSPs were evoked at 0.1 Hz for 1 hour.

### Behavioural testing

#### T-maze forced alternation task

Previously reported methods, optimised for mouse, were used to assess hippocampus-dependent learning ^[19]^. Mice were food deprived to 90% free-feeding-weight, beginning 2 days before the start of the habituation phase and with *ad libitum* access to water. Each mouse was handled at the start of food deprivation and throughout T-maze habituation for 15-20 minutes per weekday.

The T-maze was constructed from three arms, each measuring 50 x 8 cm with 10 cm colourless Perspex walls and a grey floor, mounted on a table in the centre of a room with numerous distal visual cues, such as black and white posters on the walls. Black barriers were used to block the start and goal arms. Reward consisted of a drop of Nestlé Carnation condensed milk that was placed at the end of each goal arm. Arms were cleaned with 70% ethanol between all runs to reduce odour cues. In addition, in an inaccessible well a drop of reward is always present in both arms.

Mice received 4 days of habituation to the maze, during which time they were allowed to explore the maze for 5 minutes with all arms open. During the first two days of habituation, reward was scattered along the floor and in food wells to encourage exploratory behaviour; then restricted to only the food wells at the ends of the goal arms in the last two days.

The behavioural regime lasted for five weeks, with five days of training or testing per week. Initially over the first two weeks, animals received six trials which was increased to 12 trials in the third week; each trial consisted of a sample and choice run. In the sample run, one arm was blocked off. The mouse was placed at the starting point at the base of the T, the barrier was removed and the mouse was allowed to go to the available arm and given 20 s to eat a drop of reward from the food well. For the choice run, the mouse was immediately returned to the starting point and the barrier in the previously blocked arm removed. The starting barrier was then raised and the animal allowed to choose between the two arms but only rewarded if it chose the previously unvisited arm. Thus, a correct choice was scored when the mouse selected the arm not visited in the sample run. After the choice run, the mouse was removed from the maze and placed in its holding box. The location of the sample arm (left or right) was varied pseudorandomly across the session and mice received three left and three right presentations, with no more than two consecutive trials with the same sample location. Animals were allowed a maximum of 5 minutes to make a choice to enter a goal arm in both runs before a trial was aborted. If an incorrect arm was chosen during the choice run, the mouse was confined in the arm with no reward for 20 s and then removed from the maze.

During the first three weeks of training the choice run followed immediately after the sample (test) run (there was a delay of approximately 15 s between runs for cleaning and resetting the maze). Data was analysed in blocks of 2 days and hence blocks 1-5 represent the first 2 weeks of training.

On the first day of the fourth week, mice received a repeat of the previous training sessions in order to assess retention of the task and complete the 8^th^ block. On the following two weeks, longer delays (1-10 minutes) were introduced between the sample and choice runs to extend the time that the previous choice was to be held in memory. During these intervals, each animal was placed in a separate holding box. Each mouse received two of each of the delay periods per day, varied pseudorandomly both within and across days. Squads of 15-17 mice were run per day. During training data are presented as blocks averaged across 2 days for each animal. For delays the four runs for each delay are averaged. Response times were calculated from the time that the starting block was removed until the mouse made a choice of arms and all four paws had crossed the entry point

#### Open field

The open field consisted of a plastic cylinder (diameter: 47.5 cm, height 36 cm) with a white plastic floor. Mice were placed on the periphery of the open field floor and allowed to explore freely for 30 minutes. The path of each mouse was recorded using dacQUSB recording system (Axona, St. Albans, U.K) at a sample rate of 50 Hz. The open field was optically divided into a central circle and a peripheral ring, each with equal area. The path of the mouse was analysed offline using ImageProPlus. Path length and dwell times in the periphery and centre were calculated using custom made routines written in Matlab R2010a (MathWorks).

#### Light/dark box

Analysis in the light/dark box was based on methods described by Packard et al ^[20]^. The dark box measured 20 cm x 20 cm x 30 cm, with black walls, floor and lid. The light box measured 30 cm x 30 cm x 30 cm with a white floor and light grey walls. The boxes were connected by an opening in the partition between the two compartments. An overhead light provided bright illumination in the light box. Mice were placed in the centre of the light compartment facing away from the opening and then allowed to explore for a period of 6 minutes. Time spent in each box and the number of entries into each box were recorded. An entry into a box was defined as all four paws resting inside the given box.

### Statistics

All data analysis was carried out blind to genotype. All statistics were performed using Graphpad Prism 6 with appropriately designed two-tailed t-test or ANOVA. Post hoc tests were only performed if a significant interaction between the independent variables was obtained. Animals were considered as independent samples and, where multiple data were collected from an animal, these were averaged (mean) prior to pooling. Thus sample sizes represent the number of animals. Unless stated otherwise, data are presented as mean ± SEM and differences considered significant at p<0.05.

## Results

### Identification of two separate TauD35 transgenic lines determined using TaqMan qPCR

Tau D35 express human Tau with the P301L mutation that causes hyperphosphylation of Tau ^[21]^ and was initially developed at GlaxoSmithKline as a single mouse line. However it became clear, as the genotypes were progressively uncoded in this blind study, that in some but not all types of experiments, the results from the transgenic mice fell clearly into two groups. In particular, a subset of mice showed a considerably higher density of neurofibrillary tangles at around 13 months of age than others. Moreover, in mice aged further, a subset of Tau mice showed a severely neurodegenerative phenotype, rapidly developing over about 2 weeks at 16-18 months of age, consisting of a hunched posture, piloerection and akinesia. This was so severe that it required these mice to be euthanised at this age, while others remained phenotypically normal at 24 months. We thus retrospectively determined the transgene copy number from all of the TauD35 mice included in the study.

In order to ascertain the number of transgene copies harboured by TauD35 animals, qPCR techniques were employed. Due to the similarity of the human and murine Tau sequence, transgene copy number was determined solely through the development of a probe to the CaMKIIa promoter. Assuming an endogenous murine CaMKIIa copy number of two, results gathered from transgenic animals were analysed relative to wild type copy numbers. Two clear groups within the transgenic line were apparent, one with approximately twice the copy number of the other (in the order of 10 and 5 copies respectively). Unfortunately, by the time the reason for this variability was uncovered, the HighTAU copy number had largely been bred out of the colony preventing us from increasing sample sizes in this group. The finding of this variability in copy number has the disadvantage of some experiments featuring only few animals with high copy number but the advantage is that we can at least preliminarily judge whether any effects of the TauP301L transgene are dose-dependent. Consequently, all data were pooled but with the individual copy number groups indicated in overlying scatter graphs. If strong trends or significant differences were seen between these lines then they were also analysed separately, where possible.

The two lines are referred to here as LowTAU and HighTAU, respectively. *Relative protein levels are also different in the two Tau lines, especially at younger ages* Protein levels and phosphorylation states of Tau were measured by Western blot using an array of antibodies (Fig 1). This further confirmed that the two Tau lines expressed different levels of protein in relation to their transgene copy number. Firstly, using the human-Tau-specific antibody HT-7, we assessed levels of the Tau_P301L_ protein. At 4 months of age, levels of human Tau were higher in HighTAU mice than in LowTAU mice of the same age (p=0.01; Fig 1A). This difference was less marked at 13 months but was highly significant (p<0.005). As expected, there was no immunopositive bands for human Tau observed in samples from wild type mice. The less pronounced difference at the older age is presumably because of increased Tau aggregation in the HighTAU mice which would be removed during initial centrifugation for Western blot ^[22]^. We next used the pan-Tau antibody, DA9, which, in brain tissue from wild type animals at high exposure, revealed bands at ∼50kDa, ∼58kDa and ∼64kDa, corresponding to the 3 endogenous monomeric isoforms of mouse Tau expressed in adult mice ^[23-25]^, with small increases in molecular weight due to endogenous phosphorylation, when compared to de-phosphorylated/recombinant Tau ^[26]^. The bands at the higher molecular weights are clearer when the HT-7 antibody is used to label the Sarkosyl insoluble fraction which represents the Tau tangles (Fig 1E). Moreover, although we cannot accurately quantify the protein levels in this fraction because the usual housekeeping genes are not present, we attempted to standardise the loading of protein between lanes to allow at least a qualitative comparison within ages and total protein assessed using amido black. The relatively strong signal of Tau in HighTAU mice compared to LowTAU mice supports the hypothesis that the apparently low level of Tau seen in the blots in Figs 1A-D is due to the increased deposition resulting in Tau being lost in the first stage of homogenisation which is the source of the P3 pellet from which the sarkosyl insoluble fraction (Fig 1E) is prepared.

**Figure 1.**
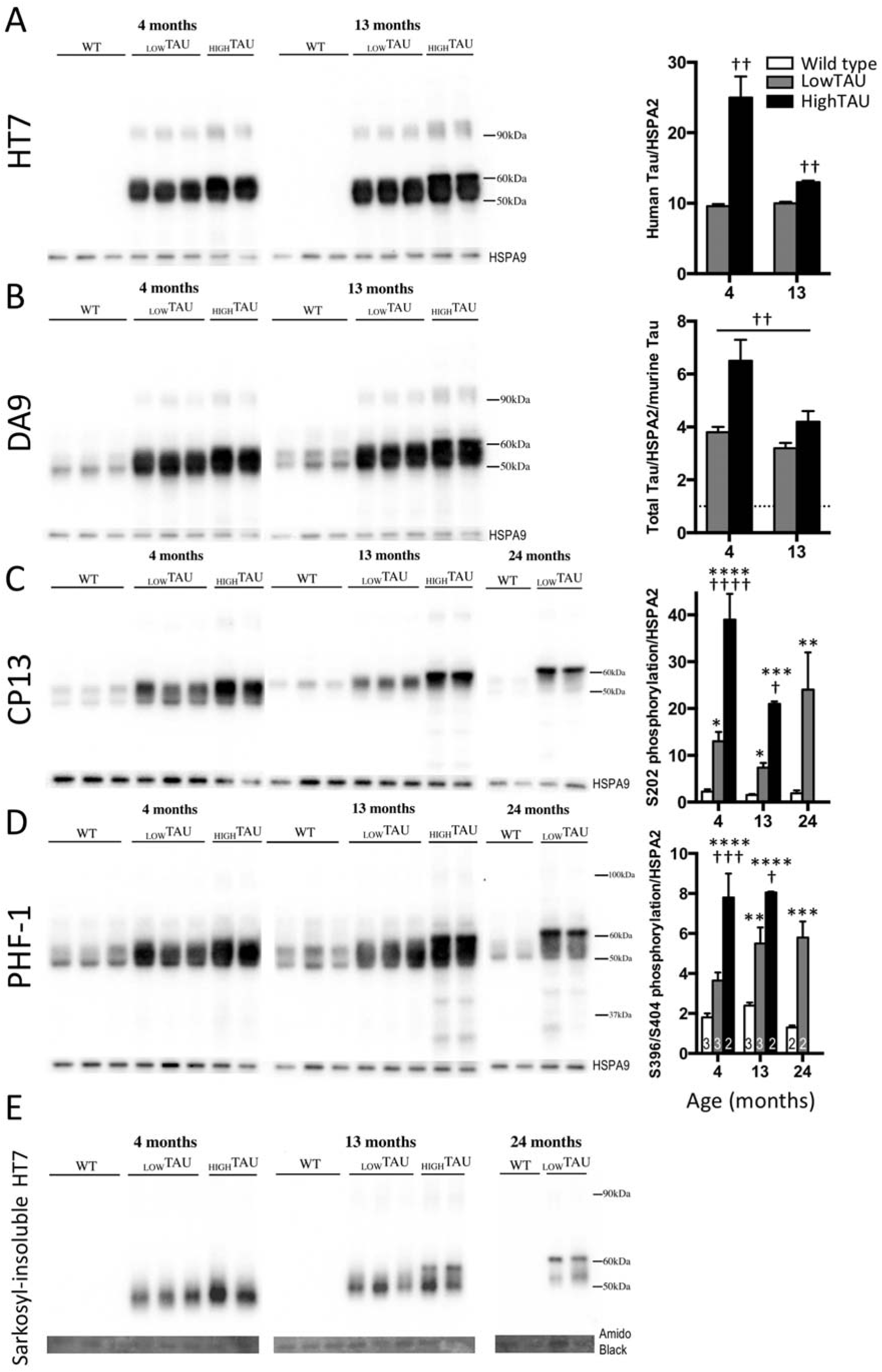
Tau protein expression and phosphorylation in TauD35 mice. Western blots were performed using antibodies specific to various Tau epitopes and quantified by normalising intensities to HSPA9. A) The human Tau-specific antibody HT7 demonstrates the TauD35 transgene is translated into protein. Two-way ANOVA genotype x age interaction, p<0.002. B) The pan-Tau antibody DA9 was used to examine total Tau (endogenous mouse and transgenic human). Values are normalised to wild type Tau levels. Note the different sized bands evident in wild type. Two-way ANOVA, main effects of genotype (p<0.003) and age (p<0.01); age x genotype interaction p=0.06; Sidak post hoc tests at 4 months: p<0.01, 13 months p>0.2). C) Levels of pS202-Tau (CP13. Age x genotype interaction: p<0.0001. D) Levels of pS396/S4040-Tau (PHF-1). Two-way ANOVA age x genotype interaction: p<0.05. E) Human Tau (HT7) within the sarkosyl-insoluble fraction. Note, Amido black (total protein) was used as a loading control, however, the low levels of protein in this fraction makes quantification impractical. Sample sizes (animals) for all probes are indicated by the numbers within columns in panel D. Sidak post hoc tests with respect to wild type are indicated by *p<0.05; **p<0.01; ***p<0.001; ****p<0.0001; and with respect to LowTAU by † p<0.05; †† p<0.01; ††† p<0.001; †††† p<0.0001.

Finally, we assessed the phosphorylation at early (S202) and late (S396/S404) sites on the Tau protein using CP13 and PHF-1 antibodies, respectively ^[13]^. A similar pattern of age-dependent Tau phosphorylation was seen with both CP13 and PHF-1, with a low level of phosphorylation present in wild type mice at all ages observed with both antibodies. Transgenic mice display a high level of both S202 (CP13) and S396/S404 (PHF-1) phosphorylated Tau when compared to wild type mice at all ages (Fig 1C&D). Interestingly, while at 4 months of age transgenic mice present phosphorylated Tau species between 50 and 60 kDa with little if any of the higher molecular weight component detectable, the larger species at around 64 kDa become clear in the 13-months old HighTAU mice but not until 24-months in the LowTAU mice, suggesting a dose and age dependent increase in this hyperphosphorylated component.

### Neurodegeneration and neurofibrillary tangle development

There were notable differences in brain size of HighTAU mice at 18 months compared to either age-matched wild type or LowTAU mice (Fig 2). This was indicated by a reduced hippocampal area, thinning of the cortex and enlargement of the dorsal ventricle. Furthermore, cresyl violet staining revealed that there was overt neuronal loss in CA1, indicated by lower cell counts at both 13 months (∼80% of wild type, p=0.07) and 18 months of age (∼40% of wild type, p<0.01; 2-way ANOVA interaction between age and genotype p=0.02; Fig 2). Similarly, there were fewer CA3 neurones in HighTAU animals, reflected by a significant main effect of genotype by 2-way ANOVA (p<0.01). In this case there was no interaction between age and genotype. We have concentrated subsequent analysis on the CA1 region but similar differences are seen throughout the hippocampus and cortex.

**Figure 2.**
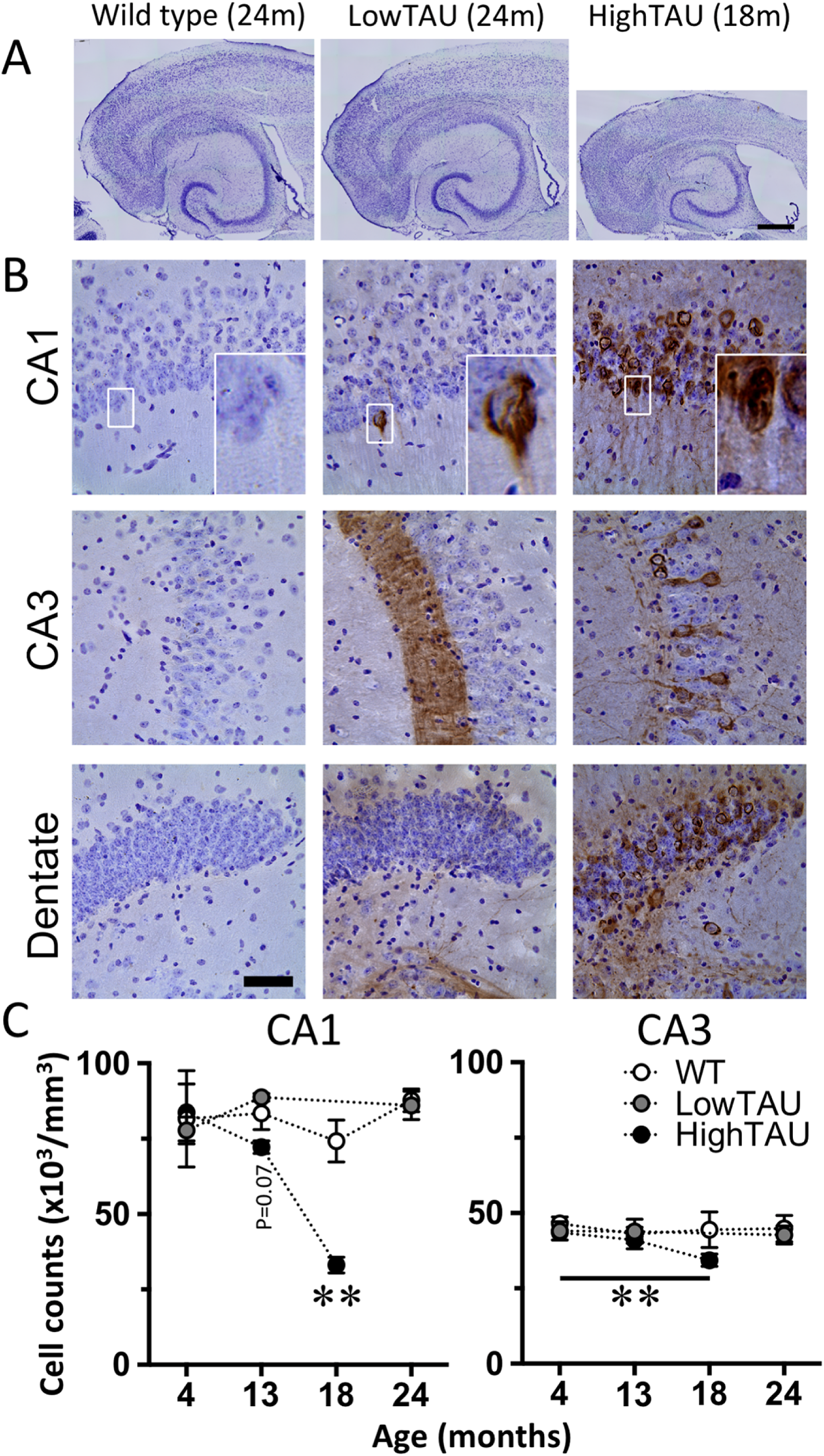
Neuronal cell loss in TauD35 mice. A) Cresyl violet stained sections of brain from 18-month-old animals showing hippocampus, dorsal ventricle and surrounding cortex. Note the distinct shrinkage of total brain tissue, an enlargement of the ventricle and thinning of the cortex in the HighTAU mice. Scale bar: 500 μm. B) Sections of primary neuronal cell body layers of CA1, CA3 and dentate gyrus stained in brown for AT8 (Tau) and counter stained with cresyl violet. Insets show mislocation of Tau to the somatic and dendritic compartments and neurofibrillary tangle-like structures within the soma in Tau mice. Scale bar: 50 μm. C) Neuronal cell counts in CA1 and CA3 indicate substantial cell loss. CA1: Two-way ANOVA age x genotype interaction for wild type v HighTAU mice, p<0.03. Sidak post hoc test **p<0.01. No significant main effects or interaction for wild type v LowTAU. (Separate two-way ANOVAs run for LowTAU v wild type and HighTAU v wild type to accommodate missing age groups). CA3: main effect of genotype, p<0.01. NB. HighTAU mice do not survive beyond 16.5-19 months of age; n=2-4 per genotype per age.

Immunohistochemistry was also carried out to assess the occurrence of neurofibrillary tangles and the regional specificity of different stages of Tau phosphorylation. Similar antibodies were employed to the Western blot experiments above, after testing specificity of antibodies in this context. Antibodies needed to be specific for Tau, recognise common pathological epitopes found in both human patients as well as other mouse models and also cover a wide range of pathological species (see methods and for review ^[13]^). To this end, a range of both commercial antibodies and non-commercial antibodies (kindly provided by Peter Davies, Albert Einstein College of Medicine, USA) were chosen, including: HT7 (human Tau), as well as AT8, CP13 (both markers for early Tau phosphorylation ^[14-15, 27]^) and MC1 (misfolded human Tau ^[16, 17]^) and PHF-1 (later Tau phosphorylation ^[15]^). A distinct pattern of immunofluorescence was observed in both CA3 and CA1 pyramidal neurones of the hippocampus, granule cells of the dentate gyrus, cells within the hilus and also the surrounding cortex (Fig 3Ai). Mossy fibre axonal projections from dentate gyrus to CA3 were particularly prominent (Fig 3Ai and Aii). The tangle-like structure of the Tau immunofluorescence were detected within the soma of neurones using confocal imaging (Fig 3Aiii).

**Figure 3.**
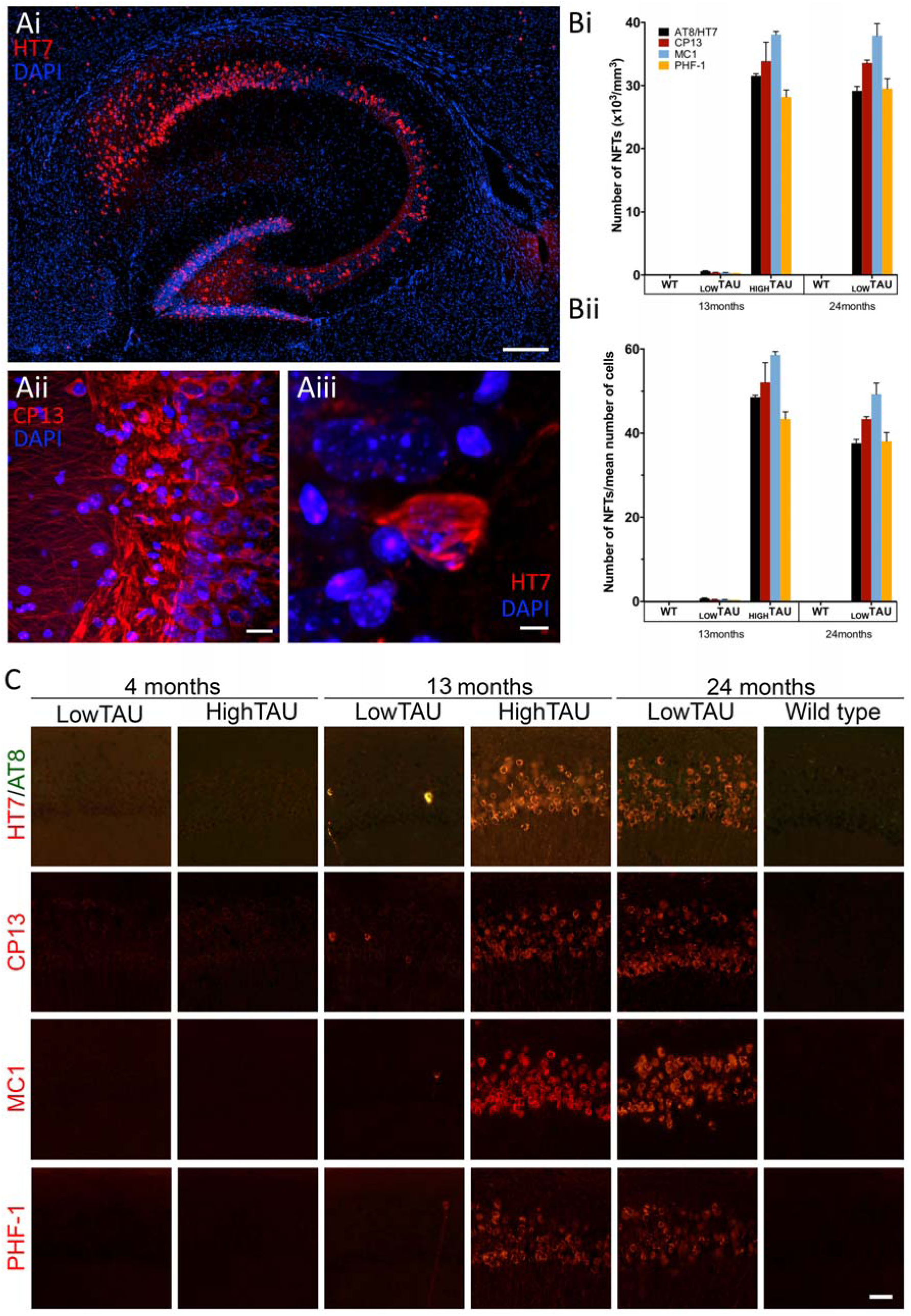
Neurofibrillary tangles in HighTAU mice. Ai) Overall image of transverse section of hippocampus from a 13-month-old HighTAU mouse, double stained using HT7 (human Tau, red) and nuclei (DAPI, blue). (Composite image using a 20x objective on an EVOS FL Auto Cell Imaging System.) Scale bar: 200 μm. Aii) CA3 region of a 4-month-old HighTAU mouse double stained using CP13 (S202) and DAPI. Note diffuse Tau staining within all cellular compartments, in particular the somatodendritic compartment. Scale bar: 20 μm. Aiii) A single CA3 neurone from Ai showing twisted Tau filaments filling the soma. Scale: 5 μm. (Images Aii and Aiii obtained using a Zeiss LSM 510 confocal microscope with 60x oil immersion objective.) Bi) Quantification of Tau immunofluorescence detected with either AT8/HT7 double staining, CP13, MC1 or PHF-1 in CA1. Bii) Data from Bi normalised to the number of neurones counted (Fig 2C). C) Example immunofluorescence images obtained from CA1 with each of the above antibodies. Scale bar: 25 μm.

Interestingly, once separated, it was clear that the 2.5-fold difference in Tau protein levels seen at 4 months between the HighTAU and LowTAU lines translated into considerable differences in the occurrence of neurofibrillary tangles by 13 months of age (Fig 3B&C). The density of tangle-like structures labelled with antibodies to all the proteins listed above by 13 months was around 100-fold higher in HighTAU than in LowTAU mice in the CA1 region (Fig3Bi). This is consistent with the effects seen above in the Western blot analysis, particularly in the Sarkosyl insoluble fraction. By 24 months, by which time neurodegeneration has resulted in the culling of all the HighTAU mice, the LowTAU mice eventually reach the high density of tangle-like staining that is seen at 13 months in the HighTAU mice (Fig 3Bi). The increase in the number of tangles remained evident when normalised to the cell density (Fig 3Bii).

### Microglia in Tau mice

We have previously reported that the same Tau mice used in the present study showed no difference in expression of *Iba1* (*Aif1*) or a range of other microglial genes at young ages, even at 8 months as neurofibrillary tangles start to appear in HighTAU mice ^[1]^. This suggested that there was little or no proliferation or activation of microglia up to this age. Here we examine distribution and phenotype of microglia in Tau mice in more detail using immunohistochemistry (Fig 4). Firstly, at 4 months of age, we confirm that there is no proliferation or activation of microglia, as measured by counting Iba1 and CD68 positive microglia, respectively, in the CA1 region in fixed sections of hippocampus in either LowTAU or HighTAU mice. By 13 months, while the wild type mice and lowTAU mice show similar age-related increases in both proliferation and activation (see also wild type mice in ^[11]^), HighTau mice show much greater changes. At this age HighTAU mice show a heavy load of neurofibrillary tangles and the start of neurodegeneration and there is a significant proliferation of microglia, (two-way ANOVA genotype x region interaction p<0.0001). This was especially strong in the *stratum lacunosum moleculare* (SLM, 3 fold change compared to WT (p<0.0001) but also with significant increases in all other layers (SO ∼2-fold increase, p<0.001; SP, SR ∼65% increase p<0.05). Moreover, activation in HighTAU mice is also increased (two-way ANOVA genotype x region interaction p<0.005) and again the increase in SLM was particularly strong (∼7 fold p<0.0001) and SO also showing a significant increase (2.7 fold, p<0.01). However, this result is difficult to interpret as the proportion of activated microglia in WT mice was very variable between individual mice, possibly relating to this age being the time when increases in activation and proliferation are occurring in all genotypes ^[11]^. Moreover, the number of HighTAU mice and the number of activated microglia per section is very low, leading to potential sampling artefacts so that only large changes could be reliably detected. Similar results were obtained in other hippocampal regions.

**Figure 4.**
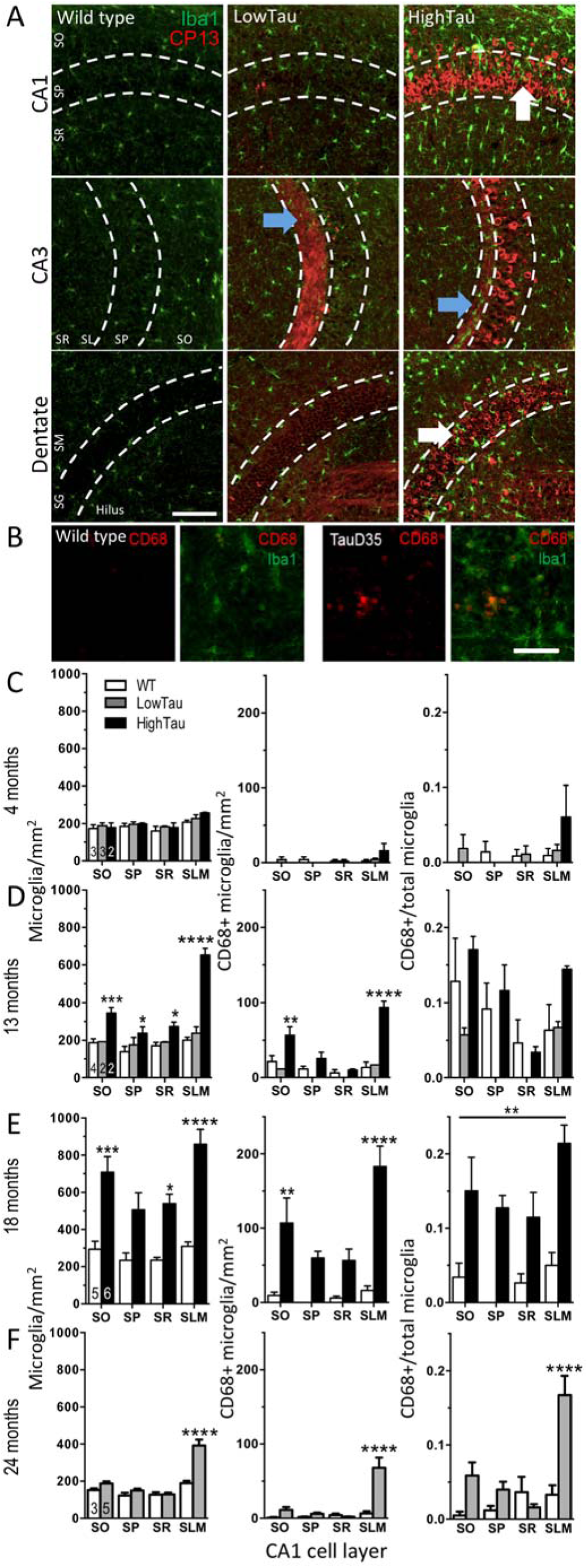
Microglia in TauD35 mice. A) Hippocampal sections showing microglia (Iba1, green) and pS202 Tau (CP13, red) from 13-month-old mice. Hippocampal layers are indicated in the wild type micrographs (SO: *stratum oriens*; SP: *stratum pyramidale*; SR: *stratum radiatum*; SL: *stratum lacunosum*; SM: *stratum moleculare*; SG: *stratum granulosum*; SLM: stratum lacunosum moleculare. Arrows indicate examples of microglia associating with Tau. B) Colocalisation of CD68 (red) and Iba1 (green) in sections from 18-month-old animals. C-F) Quantification of CA1 layer-specific alterations in 4-13-, 18- and 24-month-old mice for Iba1 positive microglia (*left*), CD68 positive microglia (i.e. where CD68 colocalised with Iba1; *centre*) and ratio of CD68 positive/total microglia (*right*). NB, in some regions, there were no CD68 positive microglia identified. At 18 months only WT vs HighTAU and 24 months only WT vs LowTAU. Sidak post hoc tests following significant layer x genotype interactions are indicated compared to wild type *p<0.05, **p<0.01, ***p<0.001, ****p<0.0001. At 18 months of age (panel E), there was a main effect of genotype for the proportion of CD68 positive microglia, **p<0.05. (NB, Sidak post hoc test for C&D were designed to also assess lowTAU versus highTAU, data not shown for simplicity). Sample sizes (animals) are indicated within columns in the Iba1 positive microglia.

Having established these initial changes, we proceeded to study the latest stages of pathology available in these mice. LowTAU mice were aged to 24 months but HighTAU mice could not be aged this far because the strong neurodegenerative phenotype that occurs at 17-18 months is considered end stage. As outlined above, at this stage there is considerable neurodegeneration. We thus compared these two groups of mice in detail, also referring back to the earlier stages described above.

The 24-month-old LowTAU mice have a similar density of tangles to the 13-month-old HighTAU mice (Fig 3) but, at this stage, they have no measurable neurodegeneration (Fig 2). Proliferation of microglia was evident at around 2-fold in the 24-month-old LowTAU mice compared to WT mice but only in the *stratum lacunosum-moleculare* and not in other CA1 regions (Fig 4F). Moreover activation (measured by CD68 positive microglia), was also increased by almost 10-fold in the SLM. Hence the SLM showed a very similar microglial response at this stage to the HighTAU mice at the equivalent stage. Interestingly there was a strong trend to show increased proliferation also in the other CA1 regions despite lack of proliferation at this stage in the 24 month old LowTAU mice suggesting that activation may precede proliferation in these mice.

In contrast, in the strongly neurodegenerating 17-18-month-old HighTAU mice, microglia both proliferate and activate across all CA1 fields (Fig 4E).

### Synaptic transmission in TauD35 mice

To study the early effects of the Tau transgene on neuronal function, whole cell voltage clamp recordings were performed. Again, retrospective genotyping was used to determine transgene copy number and data are presented as both composite mean for all Tau animals and for separated LowTAU and HighTAU mice.

Excitatory postsynaptic currents (EPSCs) were recorded from CA1 pyramidal neurones using patch clamp (Fig5Ai) and were electrically and pharmacologically isolated in the presence of the GABA_A_ receptor antagonist gabazine (6µM). While there an age-dependent increase in mean frequency of spontaneous EPSCs in cells recorded from slices of 13-month-old wild type animals, compared to those of 4-month-old animals, there was no significant difference in frequency, amplitude or decay time constant (Figs 5Aii-iv) between the genotypes at either age. In cells from the younger group, 1 μM tetrodotoxin was added to isolate action potential-independent neurotransmission miniature EPSCs. A one-way ANOVA revealed an increase in frequency in the transgenic mice (main effect of genotype p<0.05); Tukey corrected *post hoc* tests revealed a significant difference between wild type and HighTAU mice (p<0.05) and very near significance between LowTAU and HighTAU mice (p=0.053; Fig 5Aii). While there was no difference in miniature EPSC amplitudes between genotypes (Fig 5Aiii), the decay time constant was significantly different (one-way ANOVA, p<0.05) and a Tukey post hoc test revealed a significant difference between LowTAU and HighTAU mice (p<0.05; Fig 5Aiv, possibly reflecting a change in receptor subtype and/or cell geometry). Hence, while action potential-mediated activity and the resulting release of glutamate appeared to be unchanged, miniature synaptic activity was increased in the younger age group in the HighTAU mice compared to LowTAU or wild type mice.

**Figure 5.**
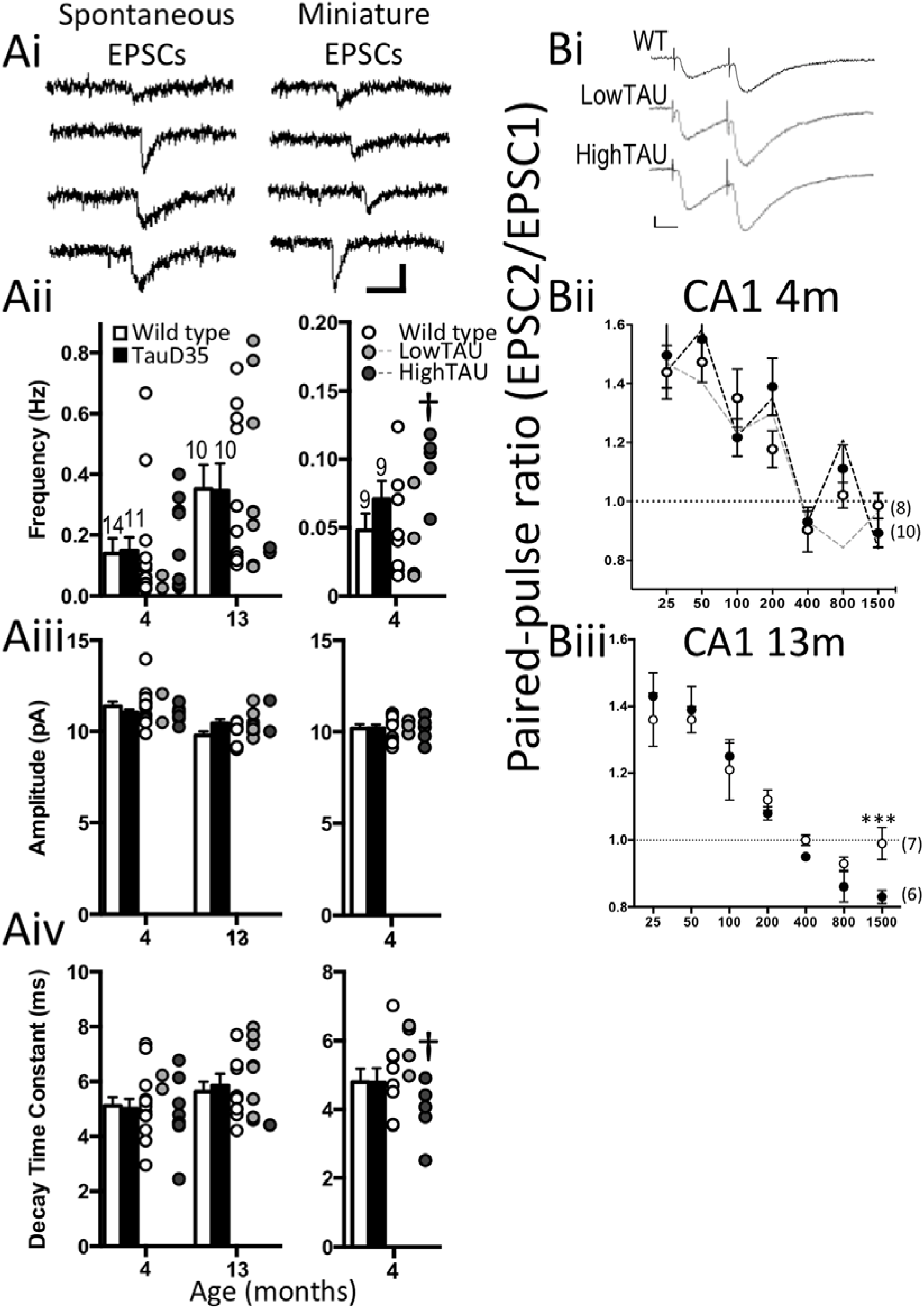
Synaptic transmission in TauD35 mice. A) Spontaneous excitatory transmission recorded from CA1 pyramidal neurones. Ai) Example spontaneous and miniature EPSCs obtained from 13-month-old animals. Scale: 10 pA x 20 ms Aii) Frequency; Aiii) Amplitude; Aiv) decay time constant (τ) for spontaneous and miniature EPSCs. Sample sizes are indicated above the bars in Aii. Individual data for LowTAU and HighTAU mice are indicated to the right of each bar plot. B) Evoked excitatory transmission. Bi) Examples of EPSCs recorded from CA1 pyramidal neurones and evoked by paired-pulses applied to Shaffer collaterals. Scale 10 pA x 20 ms. Bii & iii) Paired-pulse ratio profiles obtained from CA3-CA1 synapses of 4- and 13-month-old animals. Sample sizes are indicated in parentheses in Bii&iii. Tukey post hoc tests are indicated in panel A (*p<0.05 versus wild type; †p=0.05) following one-way ANOVA and Sidak post hoc tests are indicated in panel B following significant genotype x interval interaction by two-way ANOVA (*p<0.05; **p<0.01).

We next examined potential changes in probability of glutamate release by applying pairs of stimuli to analyse paired pulse ratios (amplitude of second response over first) from Schaffer collateral synapses onto CA1 pyramidal neurones (Fig 5Bi). As expected, synapses from wild type animals displayed paired-pulse facilitation (indicating a low probability of glutamate release), that decreased towards no change or paired-pulse depression at longer intervals. At 4 months of age there was no significant difference between transgenic mice and wild type mice regardless of copy number. At 13 months of age, post hoc genotyping revealed only LowTAU mice in this group. There were no differences in paired-pulse ratio at intervals between 25 and 800 ms, while at this age, the transgenic mice, despite having a LowTAU copy number, showed a significant depression at 1500ms compared to the wild type mice (p<0.001; Fig 5Biii). As expected wild type mice showed little if any interaction between the first and second response at this long interval. This suggests an increase in an inhibitory metabotropic autoreceptor or heterosynaptic connection in the transgenic animal.

### Magnitude of long-term potentiation is normal in Tau mice but locus of expression changes with age

Longer term synaptic plasticity was studied using extracellular field potentials recorded from *stratum radiatum* of CA1 in response to stimulation applied to the Schaffer collaterals (axons from CA3 neurones) in slices prepared from mice aged 4 to 24 months. Initially, input-output relationships were determined by applying increasing voltage pulses (Fig 6A). There were no significant differences in the field EPSP slopes recorded between the genotypes at any age. Similar to the patch clamp recordings, paired-pulse ratios did not differ between the genotypes (Fig 6B). Moreover, the magnitude of LTP (mean of responses recorded over the last 10 minutes compared to baseline) induced by a moderate tetanic stimulus did not differ between the genotypes at any age (Fig 6C&D). However, paired-pulse ratios recorded during the time course of the LTP experiment did differ. In slices from both wild type and Tau mice, the paired-pulse ratio measured at different times after induction of LTP compared to baseline showed the expected transient decrease during post tetanic potentiation (no significant difference between genotypes at any age, data not shown). While paired-pulse ratios in wild type slices returned to baseline as expected, those from Tau slices tended to remain lower than baseline (Fig 6C&E, two-way ANOVA, main effect of genotype, p<0.01). There was no significant effect of age nor an interaction between age and genotype.. This suggested that the locus of LTP induction may have a presynaptic component in the transgenic mice. As the magnitude of LTP was unchanged this may suggest a compensation between pre- and postsynaptic changes.

**Figure 6.**
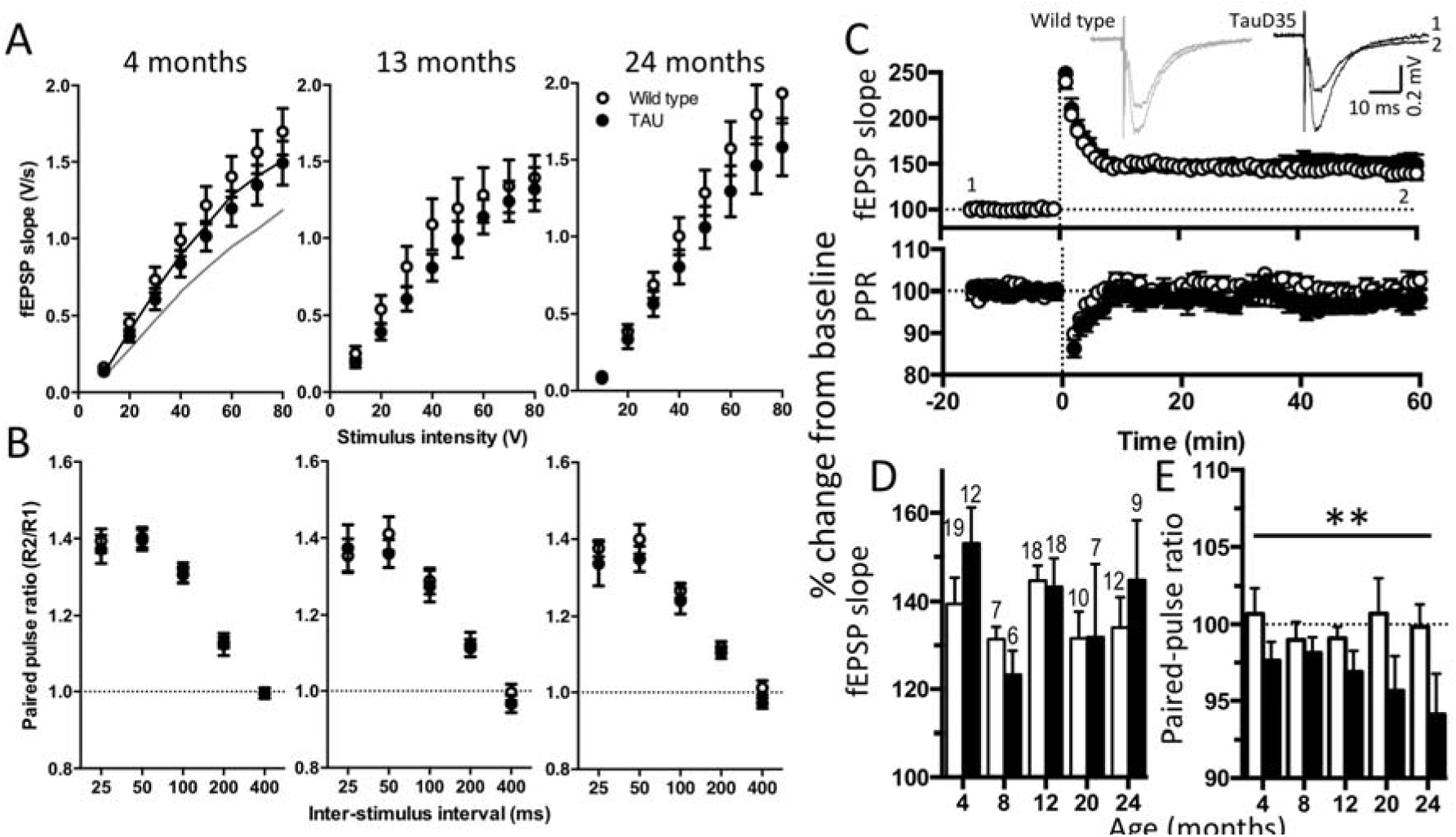
Synaptic plasticity in Tau mice. Input-output relationships at 4, 13 and 24 months of age. fEPSPs from HighTAU mice (grey line, n=8) at 4 months of age, were smaller than wild type or LowTAU mice (n=9). Paired-pulse ratios from wild type and TauD35 mice across all ages. C) LTP induced by tetanic stimulation in 4-month-old animals. Example traces from the time points indicated (1, 2) are shown above. D) Magnitude of LTP from wild type and TauD35 mice across ages. E) Change in paired-pulse ratio following induction of LTP. (Two-way ANOVA main effect of genotype but not age). Sample sizes (animals) are indicated above the bars in panel D.

### Mutant Tau mice do not show cognitive changes in hippocampus-dependent learning at 12 months of age

The hippocampus-dependent forced-alternation T-maze task was used to assess cognition in 12-month-old TauD35 mice (Fig 7A; mice used subsequently for electrophysiology or immunohistochemistry). As outlined in detail in the methods, the test consisted of a sample run followed immediately by a choice run. For the sample run one arm of the T was blocked off so that the mouse was forced to turn in one direction where they received a food reward. This was then followed by a choice run in which the both arms were open, giving them a free choice of turning left or right; the correct arm (where a food reward was then available) was the opposite arm from the sample run. Two mice, determined *post hoc* to be HighTAU mice, are reported separately from the LowTAU mice (n=9).

**Figure 7.**
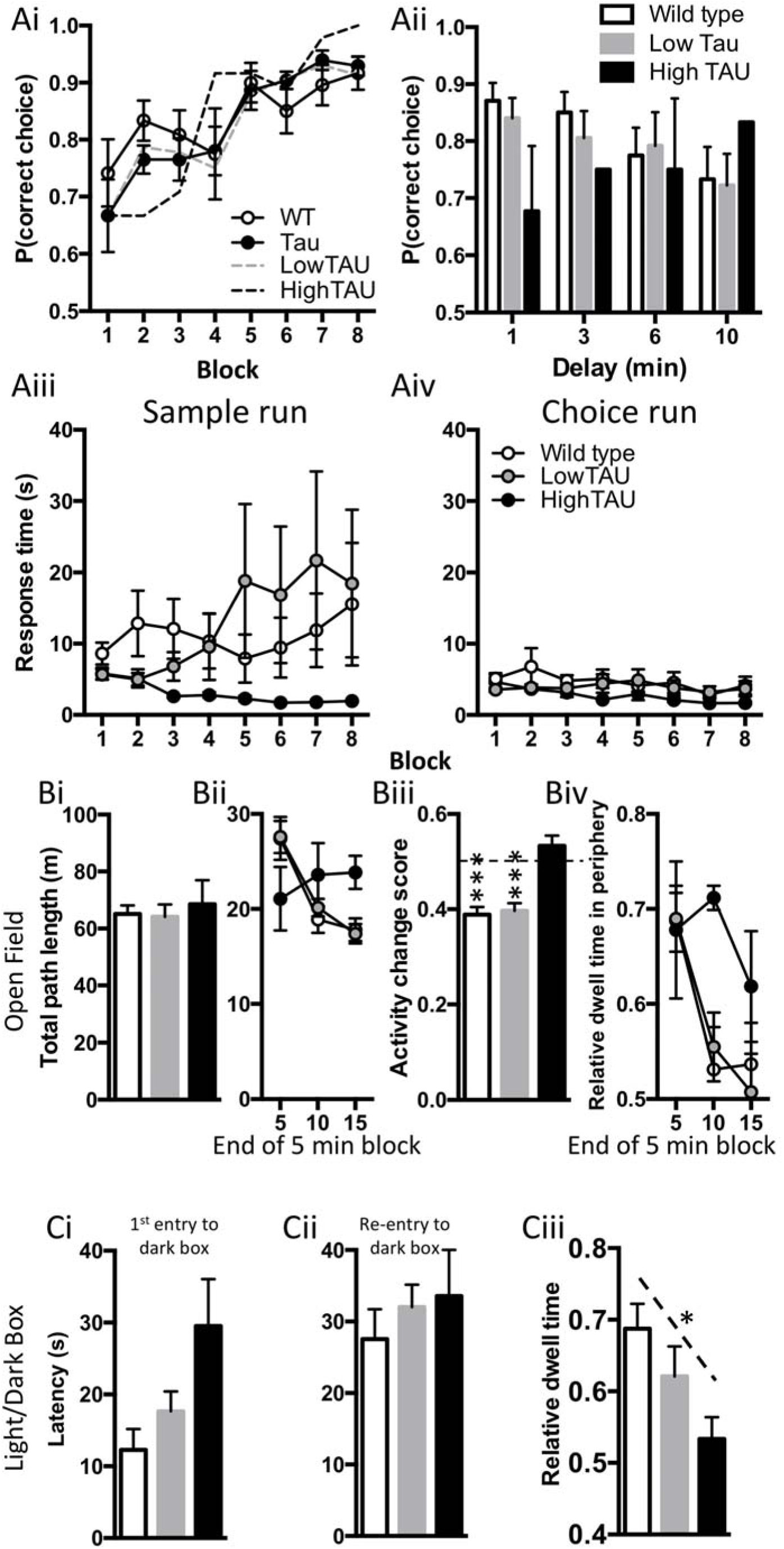
Cognitive assessment of TauD35 mice. A) Performance of mice during the training phase of the T-Maze. Wild type: n=10; LowTAU n=9; HighTAU n=2. Aii) The effect of introducing delays between sample and choice runs. Two-way ANOVA, no effect of genotype, or interaction between genotype x delay. Aiii&iv) Response times in the sample (iii) and choice (iv) runs across training blocks. B) Open field arena. Bi) Total pathlengths over the first 15 minutes of the open field exploration. Bii) Pathlengths within 5 minute periods of the open filed exploration. Biii) Habituation within the open field indicated by an activity change score [block 15/sum(block 5+block15)]. One-sample t-tests from 0.5 (i.e. no-change in activity) are denoted by ***p<0.001. Biv) Relative dwell time in the peripheral area of the open field. C) Light/Dark box. Ci) Latency to 1^st^ entry into the dark box (one-way ANOVA: p=0.06); Cii) latency to re-enter dark box; Ciii) Proportion of time spent in the dark box (NB, given the size proportions of the two boxes, 0.4 would be chance level.) There was a significant correlation between Tau transgene copy number and time spent in the dark compartment (* p=0.05).

TauD35 mice started training at similar levels to wild type mice, both improving to ∼90% over the training days (Fig 7Ai). Transgene copy number did not show any obvious differences to the outcome of the task during the training period. When the mice were challenged with delays between sample and choice runs there were no significant differences between genotypes, although surprisingly the two HighTAU mice tended to improve their performance with increased delays while both wild type and LowTAU mice showed a decrement, as expected (Fig 7Aii). However, a clear difference between the HighTAU mice and both the wild type and LowTAU mice was the rate of response in the first (sample) run of each trial (Fig 7Aiii). The wild type and LowTAU mice explored the maze, taking 5 to 20 s from being placed in the maze until all 4 feet were within the destination arm. In contrast, both HighTAU mice ran rapidly through the maze with response times starting at about 5 s but reducing to 2 s from the third block of the training period onwards. Although the sample size is small, this was consistent for both HighTAU mice in all the subsequent trials, including the delay trials, suggesting a lack of interaction with the novel environment or a behavioural disinhibition, similar to symptoms in a subgroup of frontotemporal dementia patients ^[28]^. In the choice trial there was no difference in response time between genotypes with all groups responding within about 5 seconds (Fig 7Aiv).

Unfortunately, the number of HighTAU mice was so low that no firm conclusions can be made about this potentially interesting behavioural difference but the consistency between the two mice flags an interesting point to be investigated in future studies. However, it is clear that, despite a substantial load of neurofibrillary tangles and, in the case of the HighTAU mice, a decrease in cell number, neither the HighTAU nor LowTAU mice appear to have any learning deficit in the forced T-maze.

We proceeded to test whether these possible behavioural changes reflected a change in locomotor activity or anxiety-like behaviours in the HighTAU mice (Fig 7B&C). In an open field arena, while there were no differences in the total path lengths between genotypes (Fig 7Bi), the pattern of activity was again very different for HighTAU mice. Both wild types and LowTAU mice decreased their total exploratory activity over time, while HighTAU mice maintained a constant level of exploration throughout the test period (Fig 7Bii&iii). Furthermore, the time they spent in the centre of the arena did not increase significantly over time (Fig 7Biv). Although one mouse came close to wild type levels in the final 5-minute block of the test period, the other remained largely on the edge of the arena through all trials, a behaviour that would generally be associated with anxiety.

No significant differences were seen in the light dark box (Fig 7C) when genotypes were considered as independent groups and tested by ANOVA although there was a strong trend for the Tau mice to take longer to enter the dark box (one-way ANOVA, p=0.06). Consistent with this a tendency was also observed for the transgenic mice to stay in the light for longer than the wild type mice and this was dose dependent showing a significant correlation to Tau copy number (p=0.05). In contrast to the open field result this would suggest a decrease in anxiety.

Together, these data indicate that, despite a heavy neurofibrillary tangle load and, in the case of HighTAU mice, initial neurodegeneration, there is no learning deficit measurable with the T-maze. Possible behavioural changes in habituation and anxiety particularly in the HighTAU mice would need further investigation but again would be compatible with a lack of exploratory interaction with the environment rather than specific effects of anxiety.

## Discussion

TauD35 mice express human Tau harbouring the P301L mutation under the CaMKIIa promoter, which directs expression to glutamatergic cells in the forebrain (avoiding expression in the spinal cord or peripheral nervous system). Moreover, the distribution of CaMKIIa in the brain is similar to Tau making this a very suitable promoter ^[29]^.

The P301L mutation causes frontotemporal dementia with parkinsonism linked to chromosome 17 in humans but here we were also interested in considering the role of neurofibrillary tangles in all forms of dementia, including Alzheimer’s disease. Understanding the interactions of Aβ and Tau in the development of Alzheimer’s disease is complicated by the lack of animal models that display neurofibrillary tangles and neurodegeneration dependent on rising Aβ, rather than due to mutations in Tau. While expressing mutated human Tau is useful for understanding the link between neurofibrillary tangles and neurodegeneration, it bypasses the proposed link between the Aβ and Tau proteins ^[30]^. However, this is the best approach available for studying the effects of neurofibrillary tangles on the various functional aspects of disease progression. Previous studies have described a range of mice with Tau mutations but, in general, these models have progressed rapidly, reaching neurodegeneration at relatively young ages (see Alzheimer’s Disease Research Models | Alzforum. Retrieved 14 June 2018 from https://www.alzforum.org/research-models/alzheimers-disease). In fact, a popular model, rTg4510, harbours the same mutation controlled by the same promoter but including a tetracyclineoperon–responsive element allowing suppression of the transgene. This is a rapid model that has increasing tangle load from 4 months of age and substantial neurodegeneration from about 5 months onwards ^[3]^. Moreover, spatial memory deficits are seen even earlier from around 2 months of age in these mice ^[3, 31]^. However, in humans, particularly in Alzheimer’s disease, the cognitive effects of Tau pathology and neurodegeneration are usually seen in relatively old age. Hence, the more slowly progressing model used here allows the influence of age to be considered versus the rate of pathology development. This is particularly the case when comparison of the LowTAU mice at 24 months to the HighTAU mice at 13 months at which ages their neurofibrillary tangle development is similar. What is perhaps most noticeable in these results is that, even with a heavy neurofibrillary tangle load and modest neurodegeneration seen at 13 months of age in HighTAU mice, there is little if any learning deficit, although some behavioural changes may be evident in these slowly developing models. This is similar to the human condition, in which the diagnosis of the disease only occurs after considerable brain tissue is lost ^[32]^. However, it is interesting to note that the behavioural changes seen, seem to be a lack of adaptation and interaction with the environment which may be relevant to cognitive effects in the more complex tasks of the human experience.

### Tau overexpression, phosphorylation and pathology

Phosphorylated tau is detected in both LowTAU and HighTAU animals at 4 months of age and some of human Tau is insoluble in sarkosyl. We have previously shown in TauD35 mice that rare neurofibrillary tangles could be detected in 1 out of 4 mice at 4 months and an increasing tangle load in all mice at 8 months ^[1]^. These mice have since been established to be HighTAU. In the present study aggregates are detectable by immunohistochemistry at 13 months of age in both groups. However, LowTAU animals display very few neurofibrillary tangles, along with a small decrease in the amount of phosphorylated protein compared to 4 months. In contrast, HighTAU animals display a heavy neurofibrillary tangle load at 13 months of age, corresponding to the appearance of insoluble Tau migrating at 64kDa, despite an overall decrease in phosphorylated Tau species detected by Western blot, owing to the loss of neurofibrillary tangles during homogenate sample preparation. This situation is mirrored in LowTAU animals of 24 months of age (Fig 1).

The relatively small increase in mutated human tau protein expressed in HighTAU mice (∼2.5 fold, as assessed at 4 months, Fig 1A, before substantial development of tangles) compared to their LowTAU counterparts was seen to result in a much greater difference in the timing and extent of tau pathology development. This was particularly evident at 13 months of age, when LowTAU mice were seen to display only neurofibrillary tangles within the hippocampus, compared to the extensive spread observed in HighTAU animals. At this age the ratio of neurofibrillary tangles within the CA1 region of the hippocampus in HighTAU was ∼60 times the ratio of neurofibrillary tangles calculated in LowTAU animals in the same region (Fig 2Bii). This relationship between protein level and neurofibrillary tangle development has been observed previously ^[3]^, suggesting that not only the mutation but the overall concentration of protein may be important in aggregation initiation. It is also an important observation in terms of the relevance of different overexpression models, including normal Tau protein ^[25, 33]^, in which the concentration of Tau may be a pathological factor in itself.

### Pathology and microglial proliferation

We have previously shown in mice with mutations in the Aβ pathway that the expression of microglial genes is closely related to plaque pathology, while this relationship is much weaker when compared to neurofibrillary tangle load in Tau mice ^[1]^. Here we have used immunohistochemistry to study the phosphorylation of Tau and eventual misfolding into neurofibrillary tangles in much more detail and compared this to the proliferation and activation of microglia. In the TauD35 mice, although the detection of Tau pathology is low but clear by 8 months of age in HighTAU mice, our previous gene expression study shows no change in expression of either *Iba1* or *CD68* at this age ^[1]^. Here we show with immunochemistry that also as tangles first appear in the 13-month-old LowTAU mice, that microglia are apparently unchanged but that, with the greater tangle load at this age in the HighTAU mice, proliferation is the main effect. CD68 is clearly expressed throughout the hippocampal layers once the neurodegenerative phenotype is fully developed in the HighTAU mice at 18 months of age. Increased CD68 expression is suggested to relate to an increasingly phagocytotic phenotype and appears to coincide with cell loss in these mice. This widespread microglial activation may be specifically related to the removal of damaged neurones. In contrast, at the oldest age of LowTAU mice, where no neurodegeneration is measurable, activation of microglia is also present and is particularly strong in the *stratum lacunosum moleculare*, although there is a tendency to activation in other regions. The SLM is the synaptic zone for the entorhinal cortex inputs via the temporoammonic pathway. Initially, as synapses are altered and particular axons become dysfunctional, removal of such dysfunctional axons could be protective to the ongoing network activity. This is, however, a one-way process and eventually the network would not continue effectively once substantial cell loss has occurred. Interestingly, although very delayed compared to rTg4510 mice, the neurodegenerative phenotype once it begins in HighTAU mice, is rapid. This rapidly developing neurodegenerative morbid phenotype makes cognitive testing impossible. It is interesting to note however that the lowTAU mice develop a substantial tangle load by 24 months (similar to 13 months highTAU mice) but feature a much more robust activation of the microglia at this stage but no detectable neurodegeneration. This may suggest that the microglia can protect against neurodegeneration more effectively if the Tau pathology develops more slowly.

### Synaptic transmission and plasticity

In mice with rising Aβ, we have previously reported that at 2 months in the CA1 region, even before the first plaque deposition, there is a complete loss of spontaneous action potential mediated release ^[34]^. Moreover, in evoked transmission in the Schaffer collateral pathway, an increase in glutamate release probability was observed in the Aβ mice compared to wild type mice at these early stages. These changes persisted throughout pathology development ^[11]^. In contrast, the TauD35 mice investigated here showed only very subtle changes in synaptic transmission which did not persist as tangle load developed. Perhaps not surprisingly, considering the minimal effects on basal synaptic transmission in Tau mice, LTP was not affected, although the locus of expression may have gained a presynaptic component in the Tau mice, suggesting homeostatic changes between pre- and postsynaptic compartments.

The effects on synaptic plasticity were rather more marked in Aβ mice, with an early enhancement at 2 months, followed by an almost complete loss of LTP when tetanic stimulation was used. Interestingly this was dependent on the stimulation paradigm but like the synaptic transmission changes persisted throughout pathology development. Hence, even though there is no neurodegeneration in the Aβ mice, the effects on synaptic transmission are stronger, suggesting that early physiological changes occur due to low Aβ levels in the neuropil, whereas considerable changes in Tau phosphorylation or even presence of a heavy tangle load have little effect on physiological function but rather relate directly to the death of neurones.

### Behaviour

Neither the Tau nor the Aβ mice showed cognitive deficits at 12 months of age, despite a heavy neurofibrillary tangle or plaque load respectively. At some levels this was surprising, particularly in the HighTAU mice which, unlike the LowTAU mice have substantial Tau pathology and are starting to show neuronal loss at this age. However, this is less surprising when the human disease is considered. Fox et al ^[32, 35]^ reported that people with familial genes for Alzheimer’s disease have already lost 20% of the hippocampus before they start to show symptoms, confirming that the brain is extremely good at compensating for change.

While both Tau and Aβ mice performed at similar levels as their wild type littermates in the T-maze, there were differences in motor activity and anxiety related behaviours both compared to wild type mice and comparing the different pathologies. This was reflected in both the T-maze, where HighTAU mice completed the sample run in under a quarter of the time of either wild types or LowTAU mice, and also the open field arena, where HighTAU mice failed to habituate to the arena but continued being highly active at a time when wild type mice had decreased their activity. In contrast, the Aβ mice ^[11]^ were slower to perform the T-maze task or failed to do so altogether and were less active in the open field than the wild type mice. Surprisingly, when plaque load was high at 12 months, they showed a tendency to retain memory for longer than the wild type mice.

When tests of anxiety were performed the Tau and Aβ mice initially both appeared to be more anxious, staying on the periphery of the open field. However, while this interpretation was confirmed in further tests on the Aβ mice, the interpretation of the behaviour of the Tau mice was less clear. In the light/dark box anxiety would be indicated by staying in the dark which was the result for the Aβ mice but the Tau mice stayed in the light more than wild type mice and moreover this seemed to be dose dependent in relation to copy number. In combination these behaviours suggest that there is certainly a behavioural change in the Tau mice which may indicate a lack of interaction with their environment.

The lack of memory deficits and overall subtlety of the behavioural changes reported here for both the Tau and Aβ mice, even at stages of advanced pathology but without gross neurodegeneration, is consistent with previous findings for the present Aβ model ^[36-39]^ and in other Aβ models ^[for reviews, see 40, 41]^. Deficits are seen early in the rTg4510 mouse as neurodegeneration begins ^[2-3, 31]^. The results in these more slowly developing models is however more consistent with the human condition, where, by the time sufficient deficit is evident for diagnosis, there is already a substantial reduction in hippocampal volume of up to 20% ^[32, 35]^. It is notable that in the rTg4510 mice the neurodegeneration occurs over several months (for example between 5 and 16 months ^[2]^) with behavioural testing up to 12 months ^[31]^), although this varies considerably between different studies ^[2-3, 31]^. In the much older TauD35 model reported here the initial tendency to neurodegeneration at 13 months in HighTAU mice does not result in any measurable cognitive deficit, whereas substantial neurodegeneration sets in relatively rapidly with a strong morbid phenotype developing over about 2 weeks at ages between 16.5 and 19 months preventing behavioural testing.

Given that Aβ mouse models are not a complete model of AD, i.e. lacking neurofibrillary tangle formation and neurodegeneration, which more faithfully correlate with cognitive decline, they should be considered as models of the preclinical disease, when Aβ is first deposited. The Tau models may make better models of end-stages of disease, when neurodegeneration and neurofibrillary tangle predominate the pathology. However, unless bypassed by mutations in Tau, to date, no animal model reliably offers the crucial link between rising Aβ and neurofibrillary tangles that are required to understand the full progression of the disease. It is interesting to note how well the mammalian brain can compensate and, particularly in the Tau mice how little apparent phenotype is present in terms of cognition even once neurodegeneration is under way.

## Conclusion

Given that Aβ mouse models are not a complete model of AD, i.e. lacking neurofibrillary tangle formation and neurodegeneration, which more faithfully correlate with cognitive decline, they should be considered as models of the preclinical disease, when Aβ is first deposited. The Tau models may make better models of end-stages of disease, when neurodegeneration and neurofibrillary tangle predominate the pathology. However, unless bypassed by mutations in Tau, to date, no animal model reliably offers the crucial link between rising Aβ and neurofibrillary tangles that are required to understand the full progression of the disease. It is interesting to note how well the mammalian brain can compensate and, particularly in the Tau mice, how little apparent phenotype is present in terms of cognition even once neurodegeneration is under way. The comparison of Aβ and Tau models brings up some interesting contrasts, suggesting that the levels of soluble Aβ in the general neuropil, independent of the position or even presence of plaques, has substantial effects, particularly on neurotransmitter release from glutamatergic synaptic transmission. The presence of plaques does not greatly change this, although it causes a strong immune response, possibly related to the removal of dystrophic neurites. Such dystrophic neurites only occur locally in and around the plaque, with synaptic loss having been reported to decrease with distance from the plaques ^[42, 43]^. The percentage area covered by plaques is small, even when the plaque load is heavy and hence, relative to the total number of synapses, only relatively few will be directly lost due to proximity to plaques. In contrast, although the development of Tau tangles is associated with neurodegeneration, generalised effects on synaptic responses in the wider network are more subtle until a period of rapid deterioration occurs as neurodegeneration reaches a critical stage. We suggest that this may relate to the Tau pathology causing loss of whole axons which eventually will cause network disruption in contrast to the localised effects of plaques but that in the human disease, the localised synaptic damage caused by plaques may trigger the phosphorylation and eventual dissociation of Tau from the microtubules.

## Authors’ contributions

ZJ Performed and analysed western blots, Tau immunohistochemistry, LTP and patch clamp electrophysiology and behaviour, created figures and commented on draft manuscript. PB performed LTP experiments and microglial immunohistological staining and counting at 24 months of age and commented on draft manuscript. WL performed immunohistological staining and counting of microglia at 4 and 13 months of age and commented on draft manuscript. CH and MR performed immunohistologcal staining and counting of microglia at 18 months of age and commented on draft manuscript. KY performed LTP experiments at 24 months of age. AN and ES performed LTP experiments. RD trained and supervised students for microglial staining. LM supervised behavioural experiments. RS performed genotyping and supervised students. SM designed protocol to determine Tau transgene copy number and performed genotyping. KS supervised immunohistochemistry. JCR created mice and commented on draft manuscript. FC designed and supervised behavioural experiments and commented on draft manuscript. DAS supervised molecular biology and histochemisty. DMC performed electrophysiology, supervised all experiments, wrote first draft of manuscript, created figures, performed all statistical analyses, edited and finalised manuscript. FAE obtained funding, designed and coordinated experiments, edited and finalised manuscript. All authors read and approved the final manuscript

## Acknowledgements

The authors thank Tammaryn Lashley for histological advice and the Maria Fitzgerald/Steve Hunt laboratories for use of equipment.

## References

1. Matarin M, Salih DA, Yasvoina M, et al. A genome-wide gene-expression analysis and database in transgenic mice during development of amyloid or tau pathology. Cell Rep 2015;10: 633–44. doi:10.1016/j.celrep.2014.12.041.

2. Ramsden M, Kotilinek L, Forster C, et al. Age-dependent neurofibrillary tangle formation, neuron loss, and memory impairment in a mouse model of human tauopathy (P301L). J Neurosci 2005;25: 10637–47. doi:10.1523/JNEUROSCI.3279-05.2005.

3. Santacruz K, Lewis J, Spires T, et al. Tau suppression in a neurodegenerative mouse model improves memory function. Science 2005;309: 476–81. doi:10.1126/science.1113694.

4. Yoshiyama Y, Higuchi M, Zhang B, et al. Synapse loss and microglial activation precede tangles in a P301S tauopathy mouse model. Neuron 2007;53: 337–51. doi:10.1016/j.neuron.2007.01.010.

5. Sydow A, Van der Jeugd A, Zheng F, et al. Tau-induced defects in synaptic plasticity, learning, and memory are reversible in transgenic mice after switching off the toxic Tau mutant. J Neurosci 2011;31: 2511–25. doi:10.1523/JNEUROSCI.5245-10.2011.

6. Sydow A, Van der Jeugd A, Zheng F, et al. Reversibility of Tau-related cognitive defects in a regulatable FTD mouse model. J Mol Neurosci 2011;45: 432–7. doi:10.1007/s12031-011-9604-5.

7. Spires-Jones TL, Hyman BT. The intersection of amyloid beta and tau at synapses in Alzheimer’s disease. Neuron 2014;82: 756–71. doi:10.1016/j.neuron.2014.05.004.

8. de Calignon A, Polydoro M, Suarez-Calvet M, et al. Propagation of tau pathology in a model of early Alzheimer’s disease. Neuron 2012;73: 685–97. doi:10.1016/j.neuron.2011.11.033.

9. Spires-Jones TL, de Calignon A, Matsui T, et al. In vivo imaging reveals dissociation between caspase activation and acute neuronal death in tangle-bearing neurons. J Neurosci 2008;28: 862–7. doi:10.1523/JNEUROSCI.3072-08.2008.

10. Wang JZ, Wang ZH, Tian Q. Tau hyperphosphorylation induces apoptotic escape and triggers neurodegeneration in Alzheimer’s disease. Neurosci Bull 2014;30: 359–66. doi:10.1007/s12264-013-1415-y.

11. Medawar E, Benway TA, Liu W, et al. Effects of rising amyloid beta on hippocampal synaptic transmission, microglial response and cognition in APPSwe/PSEN1M146V transgenic mice. bioRxiv 2018;420349: doi:10.1101/420349.

12. Taylor SC, Berkelman T, Yadav G, Hammond M. A defined methodology for reliable quantification of Western blot data. Mol Biotechnol 2013;55: 217–26. doi:10.1007/s12033-013-9672-6.

13. Petry FR, Pelletier J, Bretteville A, et al. Specificity of anti-tau antibodies when analyzing mice models of Alzheimer’s disease: problems and solutions. PLoS One 2014;9: e94251. doi:10.1371/journal.pone.0094251.

14. Kimura T, Ono T, Takamatsu J, et al. Sequential changes of tau-site-specific phosphorylation during development of paired helical filaments. Dementia 1996;7: 177–81.

15. Su JH, Cummings BJ, Cotman CW. Early phosphorylation of tau in Alzheimer’s disease occurs at Ser-202 and is preferentially located within neurites. Neuroreport 1994;5: 2358–62.

16. Jicha GA, Bowser R, Kazam IG, Davies P. Alz-50 and MC-1, a new monoclonal antibody raised to paired helical filaments, recognize conformational epitopes on recombinant tau. J Neurosci Res 1997;48: 128–32.

17. Weaver CL, Espinoza M, Kress Y, Davies P. Conformational change as one of the earliest alterations of tau in Alzheimer’s disease. Neurobiol Aging 2000;21: 719–27.

18. Imai Y, Ibata I, Ito D, Ohsawa K, Kohsaka S. A novel gene iba1 in the major histocompatibility complex class III region encoding an EF hand protein expressed in a monocytic lineage. Biochem Biophys Res Commun 1996;224: 855–62. doi:10.1006/bbrc.1996.1112.

19. Cacucci F, Yi M, Wills TJ, Chapman P, O’Keefe J. Place cell firing correlates with memory deficits and amyloid plaque burden in Tg2576 Alzheimer mouse model. Proc Natl Acad Sci U S A 2008;105: 7863–8. doi:10.1073/pnas.0802908105.

20. Packard MG, Introini-Collison I, McGaugh JL. Stria terminalis lesions attenuate memory enhancement produced by intracaudate nucleus injections of oxotremorine. Neurobiol Learn Mem 1996;65: 278-82. 10.1006/nlme.1996.0033.

21. Alonso Adel C, Mederlyova A, Novak M, Grundke-Iqbal I, Iqbal K. Promotion of hyperphosphorylation by frontotemporal dementia tau mutations. The Journal of biological chemistry 2004;279: 34873–81. doi:10.1074/jbc.M405131200.

22. Berger Z, Roder H, Hanna A, et al. Accumulation of pathological tau species and memory loss in a conditional model of tauopathy. J Neurosci 2007;27: 3650–62. doi:10.1523/JNEUROSCI.0587-07.2007.

23. Buee L, Bussiere T, Buee-Scherrer V, Delacourte A, Hof PR. Tau protein isoforms, phosphorylation and role in neurodegenerative disorders. Brain Res Brain Res Rev 2000;33: 95–130.

24. Liu C, Gotz J. Profiling murine tau with 0N, 1N and 2N isoform-specific antibodies in brain and peripheral organs reveals distinct subcellular localization, with the 1N isoform being enriched in the nucleus. PLoS One 2013;8: e84849. doi:10.1371/journal.pone.0084849.

25. Adams SJ, Crook RJ, Deture M, et al. Overexpression of wild-type murine tau results in progressive tauopathy and neurodegeneration. Am J Pathol 2009;175: 1598–609. doi:10.2353/ajpath.2009.090462.

26. Hanger DP, Gibb GM, de Silva R, et al. The complex relationship between soluble and insoluble tau in tauopathies revealed by efficient dephosphorylation and specific antibodies. FEBS Lett 2002;531: 538–42.

27. Bertrand J, Plouffe V, Senechal P, Leclerc N. The pattern of human tau phosphorylation is the result of priming and feedback events in primary hippocampal neurons. Neuroscience 2010;168: 323–34. doi:10.1016/j.neuroscience.2010.04.009.

28. Snowden JS, Bathgate D, Varma A, Blackshaw A, Gibbons ZC, Neary D. Distinct behavioural profiles in frontotemporal dementia and semantic dementia. J Neurol Neurosurg Psychiatry 2001;70: 323–32.

29. Lein ES, Hawrylycz MJ, Ao N, et al. Genome-wide atlas of gene expression in the adult mouse brain. Nature 2007;445: 168–76. doi:10.1038/nature05453.

30. Selkoe DJ, Hardy J. The amyloid hypothesis of Alzheimer’s disease at 25 years. EMBO Mol Med 2016;8: 595–608. doi:10.15252/emmm.201606210.

31. Blackmore T, Meftah S, Murray TK, et al. Tracking progressive pathological and functional decline in the rTg4510 mouse model of tauopathy. Alzheimers Res Ther 2017;9: 77. doi:10.1186/s13195-017-0306-2.

32. Fox NC, Warrington EK, Freeborough PA, et al. Presymptomatic hippocampal atrophy in Alzheimer’s disease. A longitudinal MRI study. Brain : a journal of neurology 1996;119 (Pt 6): 2001–7.

33. Duff K, Knight H, Refolo LM, et al. Characterization of pathology in transgenic mice over-expressing human genomic and cDNA tau transgenes. Neurobiol Dis 2000;7: 87–98. doi:10.1006/nbdi.1999.0279.

34. Cummings DM, Liu W, Portelius E, et al. First effects of rising amyloid-beta in transgenic mouse brain: synaptic transmission and gene expression. Brain : a journal of neurology 2015;138: 1992–2004. doi:10.1093/brain/awv127.

35. Ridha BH, Barnes J, Bartlett JW, et al. Tracking atrophy progression in familial Alzheimer’s disease: a serial MRI study. Lancet Neurol 2006;5: 828–34. doi:10.1016/S1474-4422(06)70550-6.

36. Pardon MC, Sarmad S, Rattray I, et al. Repeated novel cage exposure-induced improvement of early Alzheimer’s-like cognitive and amyloid changes in TASTPM mice is unrelated to changes in brain endocannabinoids levels. Neurobiol Aging 2009;30: 1099–113. doi:10.1016/j.neurobiolaging.2007.10.002.

37. Rattray I, Pitiot A, Lowe J, et al. Novel cage stress alters remote contextual fear extinction and regional T2 magnetic resonance relaxation times in TASTPM mice overexpressing amyloid. J Alzheimers Dis 2010;20: 1049–68. doi:10.3233/JAD-2010-091354.

38. Rattray I, Scullion GA, Soulby A, Kendall DA, Pardon MC. The occurrence of a deficit in contextual fear extinction in adult amyloid-over-expressing TASTPM mice is independent of the strength of conditioning but can be prevented by mild novel cage stress. Behav Brain Res 2009;200: 83–90. doi:10.1016/j.bbr.2008.12.037.

39. Howlett DR, Richardson JC, Austin A, et al. Cognitive correlates of Abeta deposition in male and female mice bearing amyloid precursor protein and presenilin-1 mutant transgenes. Brain Res 2004;1017: 130–6. doi:10.1016/j.brainres.2004.05.029.

40. Foley AM, Ammar ZM, Lee RH, Mitchell CS. Systematic review of the relationship between amyloid-beta levels and measures of transgenic mouse cognitive deficit in Alzheimer’s disease. J Alzheimers Dis 2015;44: 787–95. doi:10.3233/JAD-142208.

41. Kobayashi DT, Chen KS. Behavioral phenotypes of amyloid-based genetically modified mouse models of Alzheimer’s disease. Genes Brain Behav 2005;4: 173–96. doi:10.1111/j.1601-183X.2005.00124.x.

42. Spires-Jones TL, Meyer-Luehmann M, Osetek JD, et al. Impaired spine stability underlies plaque-related spine loss in an Alzheimer’s disease mouse model. AmJPathol 2007;171: 1304–11.

43. Kirkwood CM, Ciuchta J, Ikonomovic MD, et al. Dendritic spine density, morphology, and fibrillar actin content surrounding amyloid-beta plaques in a mouse model of amyloid-beta deposition. J Neuropathol Exp Neurol 2013;72: 791–800. doi:10.1097/NEN.0b013e31829ecc89.

